# A mathematical framework for the emergence of winners and losers in cell competition

**DOI:** 10.1101/2023.03.14.531164

**Authors:** Thomas F. Pak, Joe M. Pitt-Francis, Ruth E. Baker

## Abstract

Cell competition is a process in multicellular organisms where cells interact with their neighbours to determine a “winner” or “loser” status. The loser cells are eliminated through programmed cell death, leaving only the winner cells to populate the tissue. Cell competition is context-dependent; the same cell type can win or lose depending on the cell type it is competing against. Hence, winner/loser status is an *emergent* property. A key question in cell competition is: how do cells acquire their winner/loser status? In this paper, we propose a mathematical framework for studying the emergence of winner/loser status based on a set of quantitative criteria that distinguishes competitive from non-competitive outcomes. We apply this framework in a cell-based modelling context, to both highlight the crucial role of active cell death in cell competition and identify the factors that drive cell competition.

**Highlights:**

- Emergent winner/loser status tests for competitive outcomes in cell-based models
- Differences in biomechanical properties alone are not sufficient for cell competition
- Winners have both higher tolerance and higher emission of death signals than losers

## 1. Introduction

Cell competition is a process that occurs in multicellular organisms where cells composing genetically heterotypic tissues interact to determine their relative fitness and acquire a winner or loser status [1–6]. The loser cells are then eliminated through programmed cell death, leaving only winner cells to populate the tissue. Cell competition is context-dependent: the competing cell types are both viable in homotypic conditions, and acquire a winner/loser status only when exposed to each other in the same tissue. The main function of cell competition is to improve the overall fitness of the tissue by removing suboptimal cells. For example, during development of the *Drosophila* wing, cell competition serves as a homeostatic mechanism that stabilises tissue growth and ensures consistent wing shape [7]. It can also play a role in tumour suppression by eliminating cells with proto-oncogenic mutations [8]. However, this is not the case for all proto-oncogenic mutations: overexpression of *Myc* results in mutants that outcompete wild-type cells in a process known as super-competition [9]. This allows precancerous cells to expand within a tissue at the expense of healthy cells, without producing detectable morphological abnormalities. Cell competition can therefore also contribute to the early stages of tumour development.

The underlying mechanisms of cell competition are not yet fully understood. While progress has been made in identifying the drivers of cell competition and the pathways downstream of winner/loser identification, the intra- and intercellular processes by which cells determine winner/loser status are still unclear. Mathematical modelling, particularly cell-based modelling, has the potential to provide insight into the mechanisms of cell competition. Cell-based models allow researchers to define the behaviours of individual cells and study their effects at the population level. Because cell competition is a process that unfolds at the population level while being mediated by interactions at the cellular level, cell-based models are potentially an effective tool for exploring the most pertinent questions in cell competition. However, current cell-based models of cell competition assume *a priori* winner/loser identities [10–13]. Although such models can simulate processes occurring downstream of winner/loser identification, they do not address *how* cells become winners or losers in the first place. In this paper, we propose a mathematical framework to address precisely this question.

### 1.1. Emergence of winner/loser status

Our framework does not assume that certain cells are winners or losers *a priori*. Instead, we consider cell-based models with two cell types that vary only in their parameters and investigate the conditions that lead to competitive outcomes. Because this approach involves detecting rather than asserting winners and losers, we need a stringent definition of what a “competitive outcome” entails. We consider two defining features of cell competition: (i) both of the competing cell types are viable when grown in homotypic conditions; and (ii) the loser cells are completely eliminated in heterotypic conditions. Therefore, to identify competitive outcomes between two competing cell types in a cell-based model, we evaluate their viability in both homotypic and heterotypic conditions. This evaluation can be made either using computational simulation or through theoretical analysis, in which case viability can be analytically predicted. An interaction between two cell types is thus classified as competitive if both cell types are found to be viable in a homotypic environment and only one cell type is observed to remain viable in a heterotypic environment. These are the **cell competition criteria**, which we illustrate in Figure 1.

**Figure 1.**
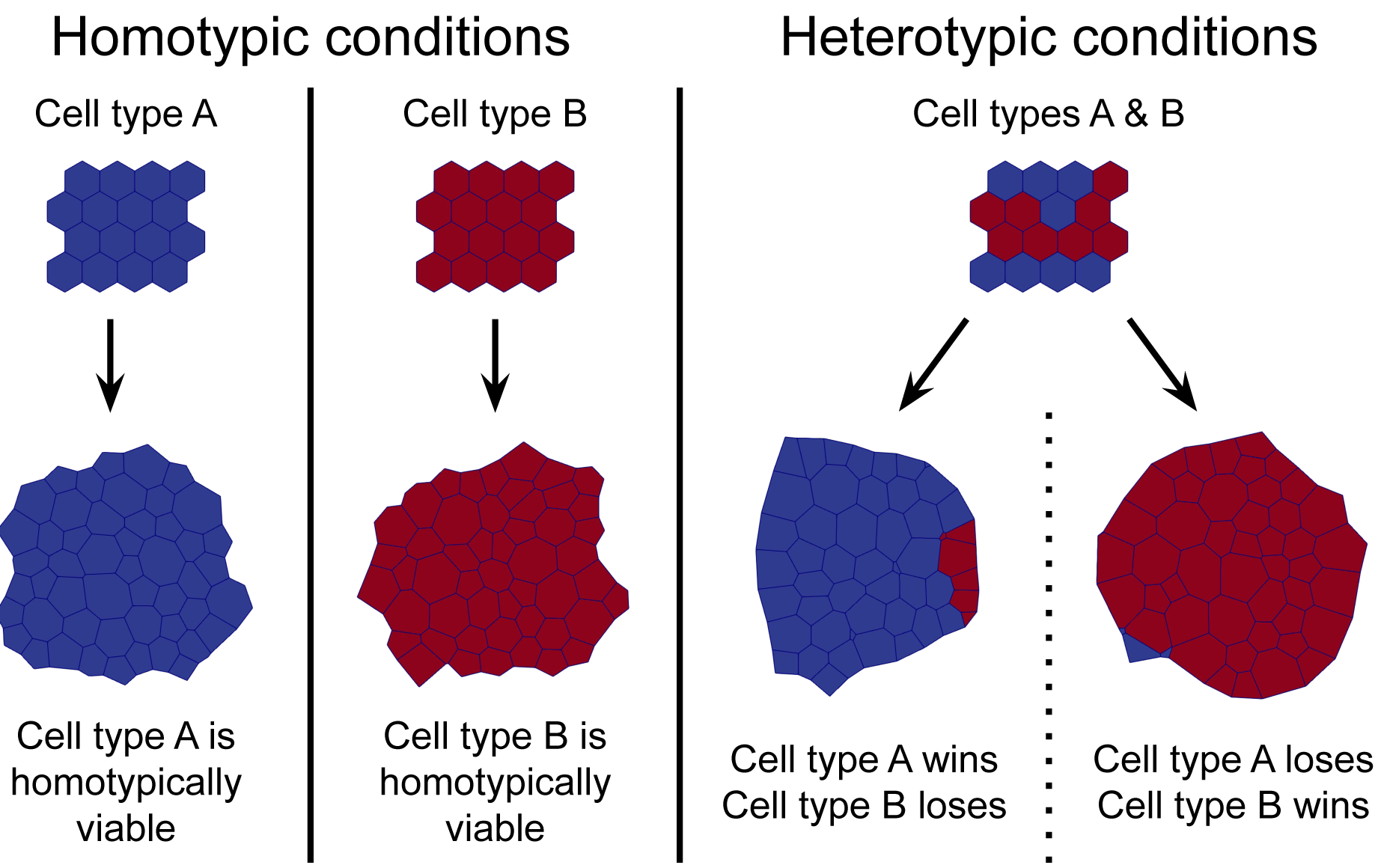
Illustration of the cell competition criteria. The two cell types A and B fulfil the cell competition criteria if (i) cell type A is homotypically viable, (ii) cell type B is homotypically viable, and (iii) cell type A is heterotypically viable and cell type B is heterotypically nonviable *or*, conversely, cell type A is heterotypically nonviable and cell type B is heterotypically viable.

We can use these cell competition criteria to identify parameter regimes that are associated with cell competition. Our approach has two important advantages over modelling frameworks that hardcode winner/loser identities. Firstly, it allows us to determine whether a given cell-based model is capable of displaying cell competition. Secondly, characterising the parameter regimes that lead to competitive outcomes helps us identify and analyse the factors that drive cell competition. Finally, we note that our framework respects the context-dependent nature of cell competition; winner/loser status is treated as an emergent property that exists only in the relationship between two cell types and is not inherent to any particular cell type.

### 1.2. Viability matrix

Generally speaking, the most appropriate definition of viability to be used for the cell competition criteria will depend on the model and the context. We assume, however, that viability is a binary property: a cell type is either viable or nonviable. Enumerating all combinations of homotypic and heterotypic viability for two competing cell types therefore results in 2^2*×*2^ = 16 possible outcomes. In order to better contextualise the cell competition criteria, we tabulate these outcomes in a **viability matrix** (Figure 2).

**Figure 2.**
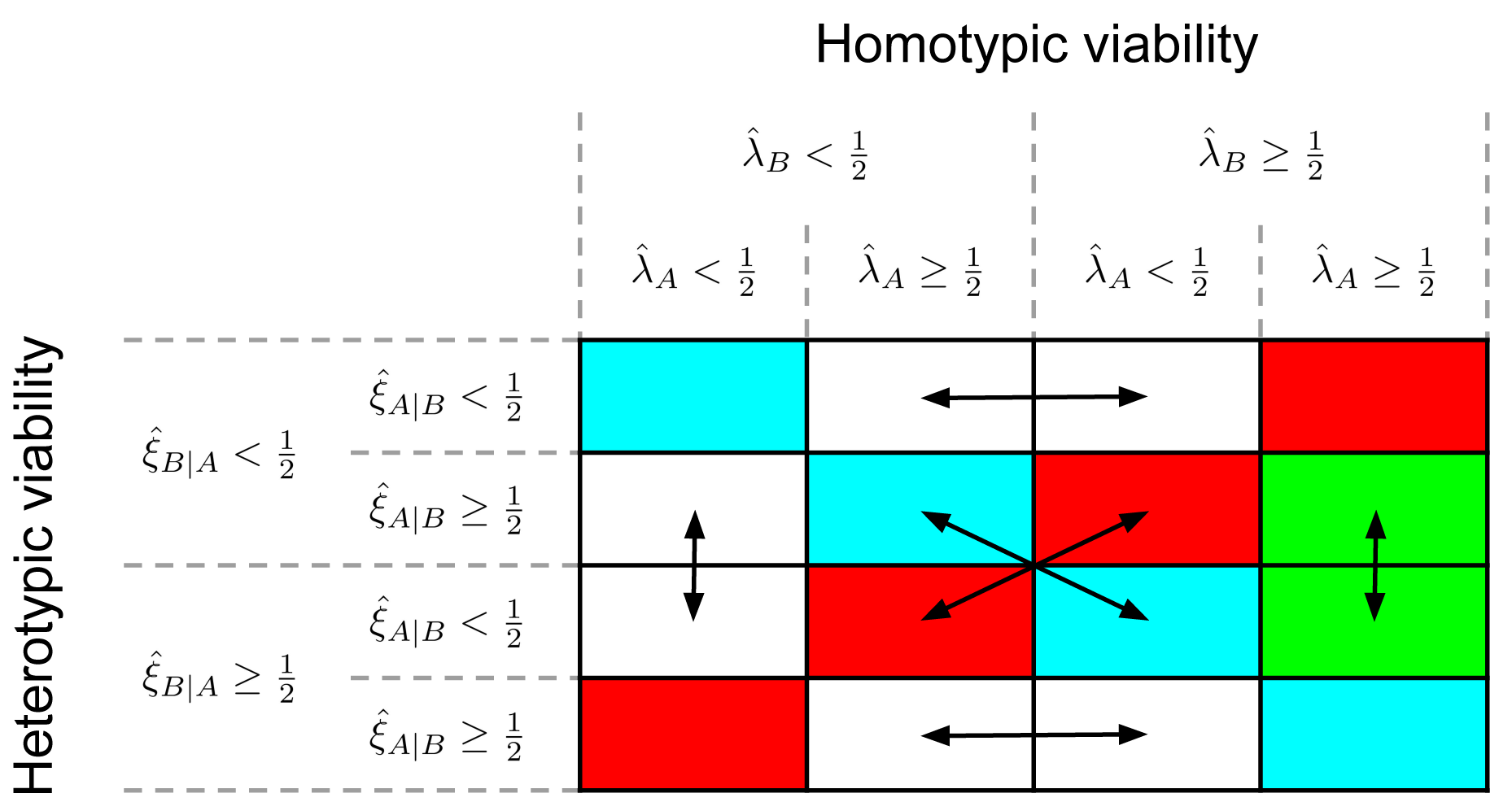
Viability matrix. The matrix is constructed by arranging the homotypic and heterotypic viability outcomes along the horizontal and vertical axes, respectively. Viability is measured in terms of survival frequency: see Equations (1) and (2) for the definitions of the homotypic survival frequencies 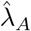 and 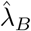, and Equations (3) and (4) for the definitions of the heterotypic survival frequencies 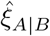 and 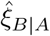 . On the main diagonal (cyan) the viability is identical for heterotypic and homotypic conditions. On the antidiagonal (red), the heterotypic viability is the opposite of the homotypic viability. The competitive outcomes are coloured green. The double-sided arrows show the result of swapping cell type labels.

In this paper, we will assess the viability of a cell population based on its *survival frequency*, which is a statistic summarising cell population growth (or decline) in simulations of cell-based models. Later on, in Section 3.3, we introduce its analytical analogue, the survival probability. Suppose, for the sake of illustration, that we have two cell types, labelled A and B, and we want to determine whether they satisfy the cell competition criteria. As Figure 1 suggests, we need to run at least two homotypic simulations, one per cell type, and one heterotypic simulation in order to measure their viability in homotypic and heterotypic conditions. We compute the homotypic survival frequencies as

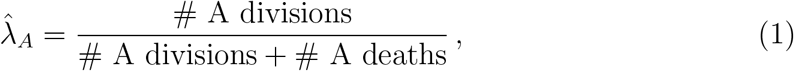

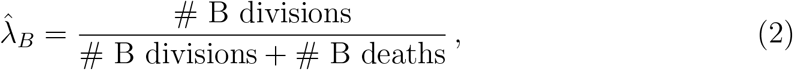

for cell types A and B from their respective homotypic simulations. Similarly, we compute the heterotypic survival frequencies from a heterotypic simulation as

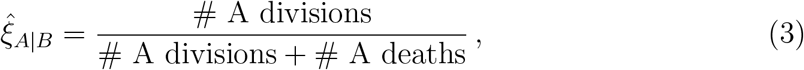

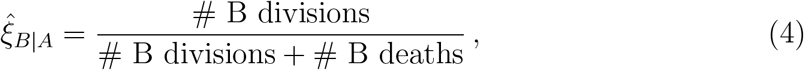

for cell types A and B, respectively. The simulations thus yield four survival frequencies: 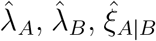, and 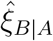 . If a survival frequency is below one half, then the cell population has declined over the course of the simulation, so we consider the population nonviable. Conversely, if a survival frequency is greater or equal to one half, then the cell population has grown or stayed the same, so we consider the population viable.

The viability matrix is then constructed by arranging the homotypic viability outcomes along the horizontal axis, and arranging the heterotypic viability outcomes along the vertical axis in the same order, as illustrated in Figure 2. Every column thus corresponds to a particular set of homotypic viability outcomes, every row corresponds to a particular set of heterotypic viability outcomes, and every element of the matrix represents a specific combination of homotypic and heterotypic viability outcomes.

The last column satisfies the first part of the cell competition criteria, i.e. both cell types are homotypically viable. Between the diagonal outcome (both cell types remain viable) and antidiagonal outcome (both cell types become nonviable) of this column, only one cell type remains viable in heterotypic conditions (green), thus completely satisfying the cell competition criteria. We define these outcomes as **competitive outcomes**. The surviving cell type is assigned the winner status, and the heterotypically nonviable cell type receives the loser status. The aim of our framework is to study the emergence of cell competition and winner/loser status by investigating the parameters and conditions that give rise to such competitive outcomes.

Finally, because we assume that the competing cell types differ only in their parameters, we note that the choice of cell type labels is arbitrary; swapping cell type labels should have no effect on the behaviour of the model. The double-sided arrows show which outcomes convert into each other as a result of swapping cell type labels, and can therefore be considered equivalent.

### 1.3. Outline

In this paper, we demonstrate the utility of the proposed framework by applying it to two different models: a mechanical model and a G2 death signal model. The mechanical model is discussed and analysed in Section 2, where we investigate whether differences in mechanical parameters between two cell types in a vertex-based model constitute a sufficient mechanism for cell competition. We perform a large parameter sweep to search for competitive outcomes, but we do not find significant evidence for competitive behaviour, suggesting that an active mechanism of cell death is necessary for cell competition. Motivated by these results, we introduce a modelling framework in Section 3 that simulates the intercellular exchange of death signals and the intra-cellular initiation of apoptosis: the “death clock” framework. Importantly, within this framework we can derive expressions for the survival probability of cells, providing us with an analytical tool for predicting the viability of cell populations. We also discuss the implementation of the death clock framework in two concrete cell-based models: the well-mixed model and the vertex-based model.

We use the death clock framework in Section 4 to construct the G2 death signal model, where cells emit death signals in the G2 phase of the cell cycle. To investigate the potential for competitive outcomes in this model, we predict the viability of cells in homotypic and heterotypic conditions using analytical arguments based on the survival probability, and validate the predictions with computational simulations of the well-mixed and vertex-based models. We demonstrate that not only can the G2 death signal model produce competitive outcomes, but also that it reveals additional biologically relevant competition regimes that have the potential to refine and expand the current theoretical understanding of cell competition. Finally, in Section 5, we discuss and interpret the results of the G2 death signal model, and propose a conceptual model of cell competition based on two key cellular properties: **tolerance** to, and **emission** of, death signals. We examine the experimental evidence in support of this model, suggest novel cell competition experiments inspired by it, and discuss potential avenues for future research.

## 2. Cell competition via differing biomechanical properties

Mechanical cell competition is a special case of cell competition, observed specifically in epithelia, that is mediated through mechanical interactions [14]. The losers in this interaction are more sensitive to cell compression than the winners and initiate apoptosis in response to cell crowding [15, 16]. In addition, we note that epithelial tissues shed live cells in response to cell crowding under homotypic conditions [17, 18]. In this study, cells undergo a “passive” form of cell death because they are extruded from the tissue as a result of mechanical interactions, and only die after being removed from the tissue. In this section, we investigate the question: are differences in biomechanical properties, combined with passive cell extrusion, sufficient to engender cell competition? A suitable cell-based framework for simulating the mechanical interactions in epithelial tissues is vertex-based modelling, since it has been shown to reproduce the dynamics of epithelial tissues in a variety of developmental processes [19].

The overall strategy of this section is therefore to construct a heterotypic vertex-based model that allows for the independent variation of mechanical parameters between two cell types, and to test whether this variation is sufficient to give rise to competitive outcomes. We call this model the “mechanical model” because our aim is to search for competitive outcomes mediated through mechanical interactions alone. In Section 2.1, we introduce the general vertex-based model and adapt it for heterotypic populations. We then describe our methodology for systematically exploring its parameter space in Section 2.2 and present the results in Section 2.3. As we will discuss in Section 2.4, we failed to find any significant evidence for competitive outcomes in the mechanical model, which motivates the construction of a model based on death signals in Section 3.

### 2.1. Vertex-based model

In vertex-based modelling, the epithelial tissue is represented by a polygonal mesh where each polygon corresponds to an epithelial cell, and the dynamics of the tissue is based on the motion of the mesh vertices. In particular, the equation of motion for vertex *i* with position **r**_*i*_, experiencing the total force **F**_*i*_, has the form [20]

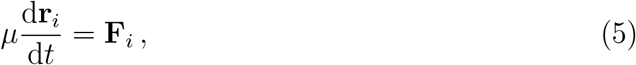

where *μ* is the friction coefficient. The force acting on vertex *i* is given by

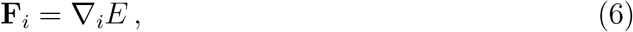

where ∇_*i*_ is the gradient of an energy function *E* with respect to the spatial coordinates of vertex *i*. We use the energy function presented in [21], which describes three major biomechanical properties: cell elasticity, cell contractility, and cell–cell adhesion;

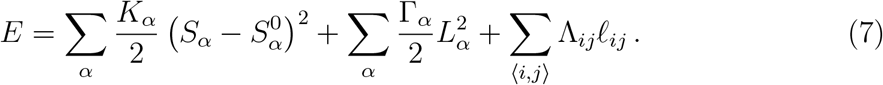

The first term represents cell elasticity, i.e. the cell’s resistance against deformation. The parameters *K*_*α*_ and 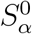 are the elasticity constant and the target cell area of cell *α*, respectively, while *S*_*α*_ is the cell area of cell *α*. The second term models cell contractility, with Γ_*α*_ and *L*_*α*_ corresponding to the contractility constant and the cell perimeter of cell *α*, respectively. The final term represents cell–cell adhesion, which is implemented as a line tension acting on cell–cell interfaces. For each edge ⟨*i, j*⟩ connecting the vertices *i* and *j*, this line tension is the product of the line tension constant, Λ_*ij*_, and the edge length, *ℓ*_*ij*_.

In addition to vertex dynamics, the vertex-based model also evolves through mesh rearrangements that allow cells to exchange neighbours, proliferate, and be extruded from the tissue. During cell division, a new edge is formed that bisects the mother cell and results in two daughter cells. Cell extrusion, on the other hand, is achieved by the “T2 swap”, which removes cells when their cell area falls below a certain threshold. There are many technical details involved with mesh rearrangements, so we refer the reader to [22] for further details.

Motivated by experiments with *in vitro* cell cultures [23], we assume a two-phase cell cycle model. The first phase corresponds to the G1 phase, and we lump together the S, G2 and M phases in the second phase. For brevity, we refer to the second phase as the G2 phase. For cell *α*, the duration of G1 phase is exponentially distributed with mean *t*_G1,*α*_. The G2 phase lasts for the fixed duration *t*_G2,*α*_. At the end of the G2 phase, cell division occurs as described above.

We divide the cell population into two non-overlapping sets that correspond to two distinct cell types, A and B. The mechanical and cell cycle constants for each cell are determined by its cell type. In particular, the elasticity constant *K*_*α*_ is given as

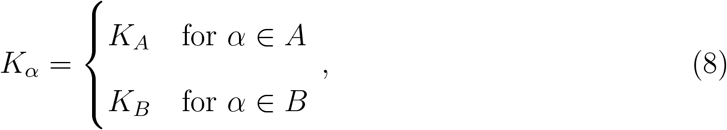

and the target cell area 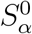, contractility constant Γ_*α*_, and cell cycle constants *t*_G1,*α*_ and *t*_G2,*α*_ are determined analogously. Since the line tension parameter is dependent on the edge type, rather than the cell type, we need to specify values for every pairing of cell types. In addition, we need to account for edges at the boundary of the tissue, which border a cell on one side and empty space on the other. Denoting the two cells sharing the edge ⟨*i, j*⟩ as *α* and *β*, we write

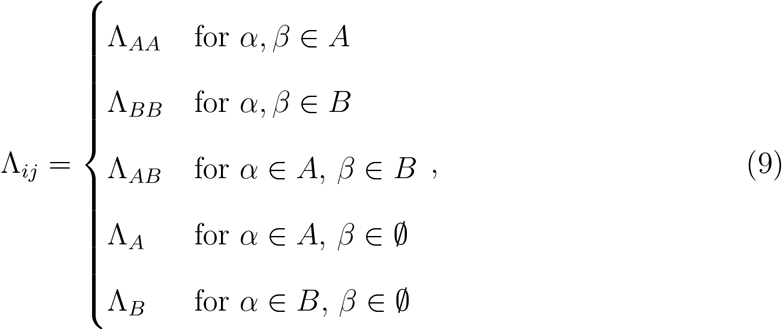

where *β* ∈ ∅ signifies that ⟨*i, j*⟩; is a boundary edge. Furthermore, we impose that each cell division results in cells that are of the same type as the mother cell, i.e. a cell of type A divides into two daughter cells of type A.

We implemented the mechanical model within Chaste, an open-source simulation package for computational physiology and biology [24] that includes a range of cell-based models [25]. We refer the reader to the following GitHub repository for the code of the mechanical model: https://github.com/ThomasPak/cell-competition.

### 2.2. Methods

After constructing the heterotypic mechanical model, we now determine whether it can generate competitive outcomes. We first performed a systematic parameter grid search varying the parameters of only one cell type, but we did not find any statistically significant evidence for competitive behaviour (results not shown). We then expanded the parameter sweep to include the parameters of both cell types. Since this involves changing the properties of two cell types simultaneously, we needed to vary twice as many parameters compared to the grid search. Therefore, because of the large number of parameters, we used a Latin hypercube sampling (LHS) method to sample parameter values. LHS methods are particularly useful when the parameter space is high-dimensional, since the number of samples required is independent of dimension [26]. In particular, we used an LHS method based on orthogonal arrays, which is an additional optimisation that improves the dispersal of parameter values [27]. Concretely, we sampled a total of 2 809 parameter sets. The lower and upper bounds for each parameter, as well as its default value, are given in Table 1.

**Table 1:** Lower and upper bounds for parameter sweep of the mechanical model. The default parameter value is also given. Any remaining parameters were set to the default Chaste values. Each simulation was given a distinct seed for generating random numbers.

| Parameter | Lower | Default | Upper |
| --- | --- | --- | --- |
| $S_A^0, S_B^0, K_A, K_B$ | 0.5 | 1.0 | 1.5 |
| $\Gamma_A, \Gamma_B$ | 0.01 | 0.04 | 0.07 |
| $\Lambda_{AA}, \Lambda_{AB}, \Lambda_{BB}$ | 0.06 | 0.12 | 0.18 |
| $t_{G1,A}, t_{G1,B}$ | 0 | 30 | 60 |
| $t_{G2,A}, t_{G2,B}$ | 40 | 70 | 100 |
| Simulation timestep |  |  | 0.05 |
| Simulation time |  |  | 250 |
| T1 threshold distance |  |  | 0.1 |
| Initial cell count |  |  | 36 |

Every parameter set thus sampled corresponds to a unique pair of cell types. For each pair, we conducted three simulations to sample the homotypic and heterotypic viabilities: two homotypic simulations (one for each cell type) and one heterotypic simulation. Each homotypic simulation has an initial population of 36 cells. For the heterotypic simulations, we split the population equally between the two cell types (18 cells each) and randomise their spatial distribution in the tissue. The homotypic and heterotypic viabilities were evaluated as described in Section 1.2, i.e. based on the homotypic survival frequency (Equations (1) and (2)) and heterotypic survival frequency (Equations (3) and (4)), respectively.

### 2.3. Results

Out of 2 809 parameter sets, 23 resulted in simulation errors because the timestep was too large. Since this only represented a tiny proportion of the parameter sweep, we excluded these parameters from our analysis. We summarised the outcomes for the remaining parameters using a viability matrix in Table 2.

**Table 2:** Count of homotypic and heterotypic viability outcomes for the parameter sweep, summarised using the viability matrix (Figure 2).

| | | $\hat{\lambda}_B < \frac{1}{2}$ | | $\hat{\lambda}_B \geq \frac{1}{2}$ | |
| --- | --- | --- | --- | --- | --- |
| | | $\hat{\lambda}_A < \frac{1}{2}$ | $\hat{\lambda}_A \geq \frac{1}{2}$ | $\hat{\lambda}_A < \frac{1}{2}$ | $\hat{\lambda}_A \geq \frac{1}{2}$ |
| $\hat{\xi}_{B\vert A} < \frac{1}{2}$ | $\hat{\xi}_{A\vert B} < \frac{1}{2}$ | 305 | 17 | 11 | 0 |
| | $\hat{\xi}_{A\vert B} \geq \frac{1}{2}$ | 0 | 407 | 0 | 4 |
| $\hat{\xi}_{B\vert A} \geq \frac{1}{2}$ | $\hat{\xi}_{A\vert B} < \frac{1}{2}$ | 0 | 0 | 476 | 16 |
| | $\hat{\xi}_{A\vert B} \geq \frac{1}{2}$ | 4 | 105 | 128 | 1 313 |

The majority of parameter sets resulted in outcomes on the main diagonal, accounting for nearly 90% of all results, indicating that little to no interaction took place in most cell type pairings. We find the second most numerous outcome in the middle entries of the bottom row, comprising 8.4% of the observed outcomes. As discussed in Section 1.2, these entries are equivalent after swapping cell labels. In these outcomes, one cell type is nonviable in homotypic conditions, but becomes viable when exposed to a homotypically viable cell type. Therefore, the most commonly observed outcomes in the parameter sweep, accounting for over 98% of all observations, are the following: heterotypic conditions either engender no changes to viability, or enhance the viability of a nonviable cell type through its interaction with a viable cell type. The latter can be construed as the opposite of a competitive outcome; the viability criteria in homotypic and heterotypic conditions are inverted with respect to the cell competition criteria.

Of the remaining categories, the largest one consists of the middle entries of the top row, accounting for 1% of observations. Similarly to the middle entries of the bottom row, only one cell type is homotypically viable. In contrast to the bottom row, however, both cell types end up nonviable in heterotypic conditions. Only 20 outcomes (roughly 0.7%) fall into the middle entries of the last column and thus fulfil the cell competition criteria, the target of our search. Finally, the least observed outcome lies on the antidiagonal (bottom left) with a total of four outcomes, or 0.1%, corresponding to the case where two homotypically nonviable cell types both become viable in heterotypic conditions.

It is important to note here that the mechanical model is stochastic, so we must account for random noise in the data. Hence, we conducted additional simulations targeting specifically those 20 parameter sets that satisfied the cell competition criteria, and tested whether the competitive behaviour was statistically significant. We found that only six out of the 20 targeted parameter sets showed statistically significant competitive behaviour with a significance level of 5%. We also ran additional simulations with segregated initial conditions to examine the influence of spatial segregation. We found that this reduced the number of significant results further to one single parameter set. We describe the methodology and results of the statistical analysis in more detail in Section S1 of the supplementary material.

### 2.4. Discussion

In this section, we constructed a heterotypic vertex-based model, namely the mechanical model, to investigate whether differences in mechanical properties are sufficient to give rise to cell competition. We performed a large parameter sweep and found that we could only reliably reproduce competitive behaviour for a tiny fraction of the simulated parameter sets. Most of the parameter sets resulted in no observable interactions, and most of the interactions that did occur generated the opposite outcome of cell competition.

We conclude that simply varying the parameters of the mechanical model is not sufficient to reliably generate competitive behaviour. This agrees with experiments suggesting that cell competition generally depends on an active mechanism of cell death, such as apoptosis [28], and that mechanical cell competition is no exception in this respect [15, 16]. We note that these results do not exclude the possibility of mechanical interactions playing a role in cell competition. They do strongly suggest, however, that passive cell death alone is an insufficient mechanism for cell competition and that mechanical interactions must be paired with an active mechanism of cell death to produce robust competitive behaviour.

## 3. Cell competition via exchange of death signals

The results of Section 2 suggest that cell competition requires an **active** and **non-autonomous** mechanism of cell death. This observation is also supported experimentally [7, 8, 28]. Therefore, the aim in this section is to develop a modelling framework for cell competition implementing such an active and non-autonomous mechanism for cell death. The core idea is that cells exchange “death signals” with their neighbours and that these signals are accumulated by the cell into an abstract quantity called the “death clock”. When the death clock reaches a threshold value, apoptosis is triggered. We do not yet attach the death signal to a concrete biological mechanism because there are multiple competing hypotheses regarding the mode of intercellular communication that underlies cell competition, and because the mode of communication may depend on the specific type of cell competition under consideration.

We first discuss our biological assumptions and modelling choices in Section 3.1, before introducing the death clock framework in Section 3.2. In Section 3.3, we define the survival probability and derive its analytic expression for a given death signal. Crucially, the survival probability enables us to analyse the death clock framework from a theoretical perspective and make predictions on the viability of cell populations. Finally, in Section 3.4 we discuss the implementation of the death clock framework in two computational cell-based models: the well-mixed model and the vertex-based model. The analytical and computational tools presented in this section will be used in Section 4 to conduct a thorough investigation of the G2 death signal model.

### 3.1. Assumptions

A series of studies involving mathematical modelling and experiments have revealed the importance of threshold mechanisms in the initiation of apoptosis [29–32]. For instance, it was shown that death ligand-induced apoptosis requires a threshold proportion of ligand to receptor numbers to be reached [31, 32]. Given this precedent, we propose a model in which competition-induced apoptosis is triggered by the accumulation of death signals reaching a threshold value.

Furthermore, it has been established in the literature that apoptosis and the cell cycle are closely coupled [33–36]. Notably, the regulatory protein *Myc* is known to affect both cell cycle progression and apoptosis [37–39]. On the one hand, *Myc* is necessary for the transition of G1 to S phase, and it induces cell cycle progression in quiescent cells [37, 39]. On the other hand, *Myc* has been associated with increased rates of cell death [38]. Coupled with the fact that differential *Myc* expression results in cell competition [9], we hypothesise that apoptosis, competition, and the cell cycle are interrelated. Concretely, we assume that the cell is only susceptible to competition-induced apoptosis in G1 phase, and that the cell is committed to division from S phase onwards. Similar to the vertex-based model in Section 2, we assume a two-phase cell cycle model, where we treat the duration of the G1 phase as a random variable and lump together the S, G2 and M phases into the G2 phase, which has a fixed duration.

### 3.2. Death clock framework

The death clock framework consists of two coupled cellular processes: the cell cycle and the death clock, where the death clock governs the initiation of apoptosis in response to death signals. We consider the cell cycle to be an **autonomous** process, meaning that it is not affected by other cells. On the other hand, the death clock is a **non-autonomous** process because it is driven by extracellular signals produced by other cells. Together, these processes determine whether and when the cell divides or initiates apoptosis.

At division, we sample a stochastic G1 duration, denoted as *t*^∗^, from the **G1 duration distribution** *𝒞*, i.e.

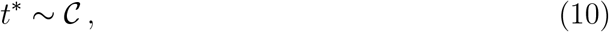

where *𝒞* is subject to the constraints that (i) *t*^∗^ ∈ [0, ∞) and (ii) *E*(*t*^∗^) = *t*_G1_, with *t*_G1_ the **autonomous G1 duration**. If apoptosis is not triggered by the death clock, the cell spends a duration *t*^∗^ in G1 phase and then transitions into G2 phase. After spending a fixed duration, *t*_G2_, in G2 phase, the cell divides and the process repeats for each of the daughter cells.

We model the accumulation of death signals using an ordinary differential equation (ODE) model in which the **death clock**, denoted by *τ* (*t*), evolves according to the ODE

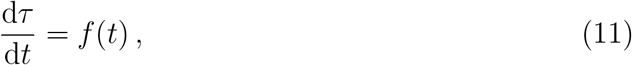

where *f* (*t*) ≥ 0 is the **death signal** experienced by the cell. At birth, the death clock of a cell is initialised to zero, i.e. *τ* (*t* = 0) = 0. The apoptosis rule is then

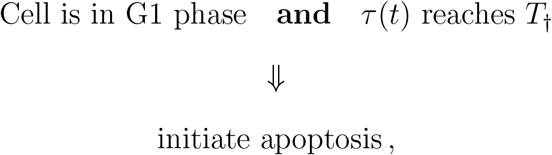

where *T*_†_ is the **death threshold**. We define the **survival condition** as

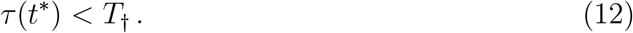

We note that there are two potential sources of uncertainty in the death clock framework: variability in G1 duration and in the death signal. The former originates from the cell cycle, the latter from intercellular interactions, and both contribute to the decision of the cell to initiate apoptosis. Our framework can thus be regarded as a minimalist model of autonomous and non-autonomous processes interacting to govern competition-induced apoptosis. The death clock framework is summarised by the flowchart in Figure 3.

**Figure 3.**
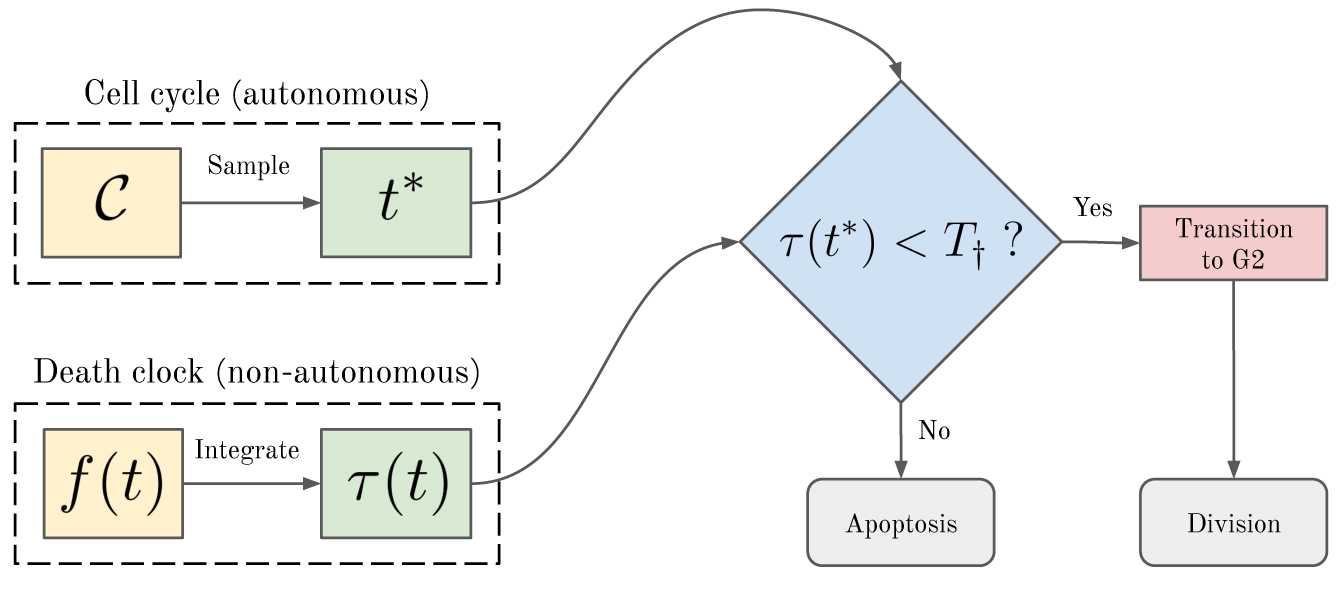
Death clock flowchart. The “Sample” step corresponds to Equation (10), and the “Integrate” step corresponds to Equation (11). The condition in the decision block is the survival condition, corresponding to Equation (12).

### 3.3. Survival probability

In order to predict the viability of a cell population, we must determine the probability of cells surviving. This problem is intractable when considering the uncertainty in the death signal and in the cell cycle simultaneously. To make analytic progress, we fix the death signal and consider exclusively the variance in the cell cycle, which lets us derive an expression for the “survival probability”. We define this survival probability, which we denote by *θ*, as the probability that the survival condition (Equation (12)) is satisfied, i.e.

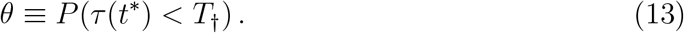

Assuming that *f* (*t*) is a non-negative integrable function, we define

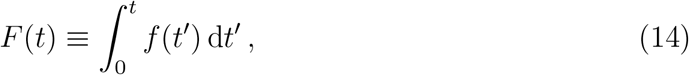

such that the value of the death clock at time *t*^∗^ is *F* (*t*^∗^). This lets us write the survival condition as *F* (*t*^∗^) *< T*_†_. We define the pseudoinverse function of *F* (*t*) as

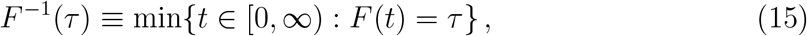

so that we can reformulate the survival condition as

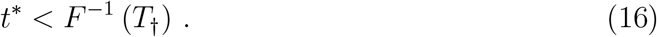

Substituting this into Equation (13), and denoting the cumulative distribution function for the distribution of *t*^∗^ as Ψ(*t*), we obtain

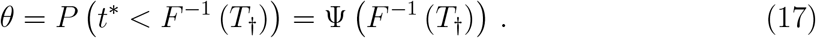

As a special case, consider the constant death signal *f* (*t*) = *c*, where *c >* 0 is a positive constant. We then have *F* (*t*) = *ct* ⇒ *F* ^−1^(*τ*) = *τ/c* ⇒ *θ* = Ψ(*T*_†_*/c*).

### 3.4. Cell-based death clock models

So far, we have described the processes leading to competition-induced apoptosis from the perspective of a single cell. The death clock framework can be embedded in any cell-based model that (i) provides cells with an extracellular environment from which to derive a death signal and (ii) includes a cellular operation for initiating apoptosis. In this paper, we implement the death clock mechanism in two particular cell-based models: the vertex-based model (Section 2.1), and a well-mixed model. In the vertex-based model, a cell interacts only with cells in its local neighbourhood. In the well-mixed model, on the other hand, each cell interacts with all other cells on an equal basis. In Section 4, we use both models in a complementary manner. Here, we present a high-level outline of the well-mixed and vertex-based models. For a detailed discussion of their numerical implementations, see Sections S2 and S3 in the supplementary material. We provide the code for both models in the following GitHub repository: https://github.com/ThomasPak/cell-competition.

#### 3.4.1. Well-mixed model

For each cell *α*, we represent its state with the cell vector **y**_*α*_(*t*), where *α* = 1, …, *N* (*t*), and *N* (*t*) is the number of cells at time *t*. We write the cell vector as

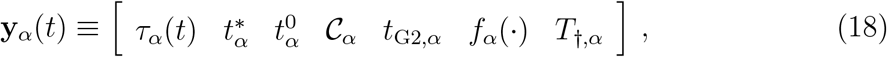

and summarise its contents in Table 3. The state of the system, denoted *S*(*t*), is then

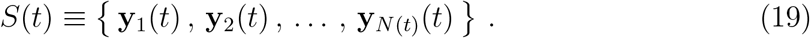

We evolve the death clock for each cell *α* as

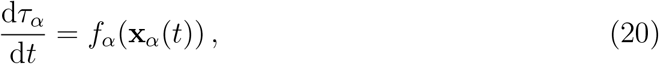

where *f*_*α*_(·) is the death signal function and **x**_*α*_(*t*) is the “input vector” representing the extracellular environment. Since the cell population is well-mixed, this environment is composed of every cell except itself, i.e. **x**_*α*_(*t*) = [**y**_1_(*t*), …, **y**_*α*−1_(*t*), **y**_*α*+1_(*t*), …, **y**_*N*(*t*)_(*t*)] .

**Table 3:** Summary of cell vector elements.

| Symbol | Description |
| --- | --- |
| $\tau_\alpha(t)$ | Death clock |
| $t_\alpha^*$ | Sampled G1 duration |
| $t_\alpha^0$ | Birth time |
| $\mathcal{C}_\alpha$ | G1 duration distribution |
| $t_{G2,\alpha}$ | G2 duration |
| $f_\alpha(\cdot)$ | Death signal function |
| $T_{\dagger,\alpha}$ | Death threshold |

In addition, we define two discrete operations: cell division and cell death. When a cell’s age reaches its total cell cycle duration, the division operation is triggered which constructs two daughter cells; one in a new cell vector and one reusing the mother cell vector. When a cell’s death clock reaches the death threshold in G1 phase, the cell is removed from the population. See Section S2 in the supplementary material for further implementation details.

#### 3.4.2. Vertex-based model

We implemented the vertex-based death clock model by augmenting the basic vertex-based model, introduced in Section 2, with the death clock mechanism. Briefly, this involves equipping every cell with a death clock that can trigger apoptosis. The death clock for each cell is evolved similarly to the well-mixed model using Equation (20). However, the input vector **x**_*α*_(*t*) is constrained to contain only information about the local extracellular environment of cell *α*, for instance the states of its direct neighbours. Apoptosis is implemented in the vertex-based model by shrinking the target cell area, 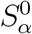, to zero, which causes the cell to contract until it is extruded from the tissue. See Section S3 in the supplementary material for implementation details.

## 4. The G2 death signal model

Having introduced the death clock framework, as well as the analytical and computational tools to investigate its dynamics, we now turn our attention to a particular form of the death signal, namely the G2 death signal. In the G2 death signal model, cells emit death signals to their neighbours while they are in G2 phase. This choice is motivated by the observation that cell competition often manifests as patches of proliferating cells inducing apoptosis in neighbouring cells to make room for themselves. In the death clock framework, cells in G2 phase are committed to division, so we decided to associate the death signal with the decision to proliferate. Moreover, experimental evidence suggests a link between cell cycle progression and death signals [40, 41]. Concretely, the G2 death signal is defined as

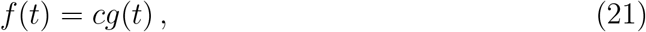

where *g*(*t*) is the proportion of neighbouring cells in G2 phase, i.e.

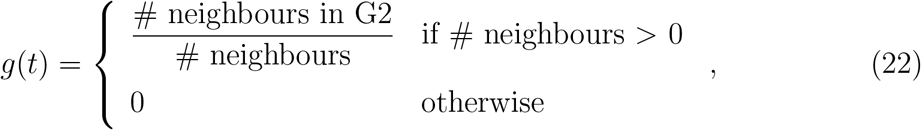

and *c* is a positive constant.

We first investigate the effect that the G2 death signal model has on homotypic populations in Section 4.1. This is done by deriving an expression for the homotypic survival probability (Section 4.1.1), which further enables us to characterise the parameter space in terms of homotypic viability (Section 4.1.2). For heterotypic populations (Section 4.2), we similarly characterise the heterotypic survival probability (Section 4.2.1) and use it to derive the conditions for viability in each subpopulation (Section 4.2.5).

We also describe and classify the different types of competitive interactions encountered in the G2 death signal model in Section 4.2.4.

The cell competition criteria are based on both the homotypic and heterotypic viabilities, so the results of Sections 4.1 and 4.2 are combined in Section 4.3 to identify biologically relevant competition regimes. Notably, we demonstrate that the G2 death signal model is capable of producing competitive outcomes. Furthermore, our detailed investigation of the parameter space reveals additional competition regimes that refine and generalise the classical competition regimes defined in the literature. Finally, in Section 5 we provide a detailed discussion of our findings and their implications for cell competition.

### 4.1. Homotypic populations

We defined the survival probability in Section 3.3 for a given death signal, but in general the death signal received by any particular cell is not known *a priori*. Fortunately, as we will see in Section 4.1.1, we can derive a useful approximation of the death signal in the G2 death signal model and use this to characterise the homotypic survival probability. In Section 4.1.2, we build on this result to characterise the *proliferation regimes*, which we define as the parameter regimes in which cells are viable or nonviable. Finally, we validate these proliferation regimes using simulations of the well-mixed and vertex-based models in Section 4.1.3.

#### 4.1.1. Homotypic survival probability

In order to derive the homotypic survival probability, we need to obtain an expression for the G2 death signal under homotypic conditions. But first, we highlight the critical role of the cell cycle in the G2 death signal model to motivate the definition of an important dimensionless parameter.

In the G2 death signal model, cells only emit death signals in G2 phase and this leads to an important trade-off; cells in G1 phase are vulnerable to death signals and do not generate death signals, whereas cells in G2 phase are impervious to death signals but do generate death signals. This raises the question: what is the impact of changing the proportion of the cell cycle that is spent in G1 or G2 phase on the survival probability, given a fixed total cell cycle duration? In order to investigate this question, we denote the total cell cycle duration as *t*_G_, and define *β* as the fraction of the cell cycle that is spent, on average, in G1 phase, so that

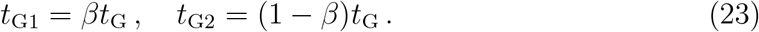

Even though cell cycle phases are stochastic in the G2 death signal model, we found that the death signal is not only relatively stable, but also predictable. In particular, we observe that the system is **ergodic**, in the sense that the average proportion of cells in G2 phase relative to the population well approximates the average proportion of the cell cycle spent in G2 phase. More precisely, we state that the system is ergodic if, on average,

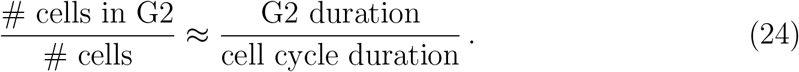

Furthermore, if the system is well-mixed, then we can approximate *g*(*t*) as

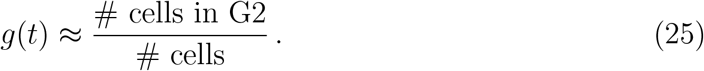

Combining Equations (24) and (25), we have

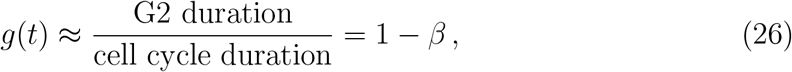

so that the death signal is

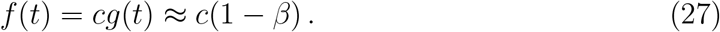

Applying the methodology of Section 3.3, we use this result to derive the **homotypic survival probability**, denoted *λ*, as

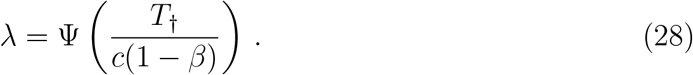

For an exponential cell cycle model more specifically, this becomes

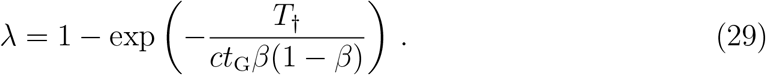

In order to simplify the notation, we introduce the dimensionless parameter *η*,

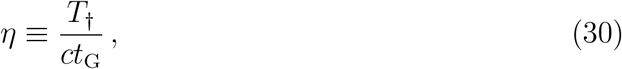

which can be interpreted as a normalised death threshold. Hence, we write the homotypic survival probability as a function of two dimensionless parameters:

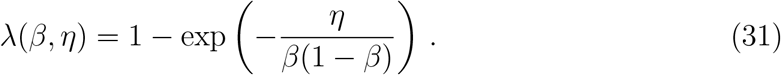

We validate this expression via simulation in Section S4 of the supplementary material.

#### 4.1.2. Homotypic proliferation regimes

Based on the homotypic survival probability *λ*, we distinguish between two proliferation regimes for homotypic populations^1^:

##### Nonviable Regime 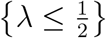

Cells are equally or more likely to die than to proliferate, hence the population declines. We say that cell types in this regime are **nonviable**.

##### Viable Regime 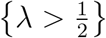

Cells are more likely to proliferate than to die, hence the population grows. We say that cell types in this regime are **viable**.

We define the **homotypic viability curve** as the curve satisfying *λ* = 1*/*2. This curve separates the Nonviable Regime from the Viable Regime. For the exponential cell cycle model, the homotypic viability curve is given by

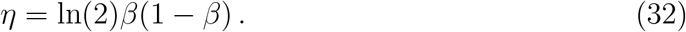

This analysis therefore predicts that a population is viable for all *η >* ln(2)*/*4, and for *η* ≤ ln(2)*/*4 it is viable for extreme values of *β* and nonviable otherwise (Figure 4(a)).

**Figure 4.**
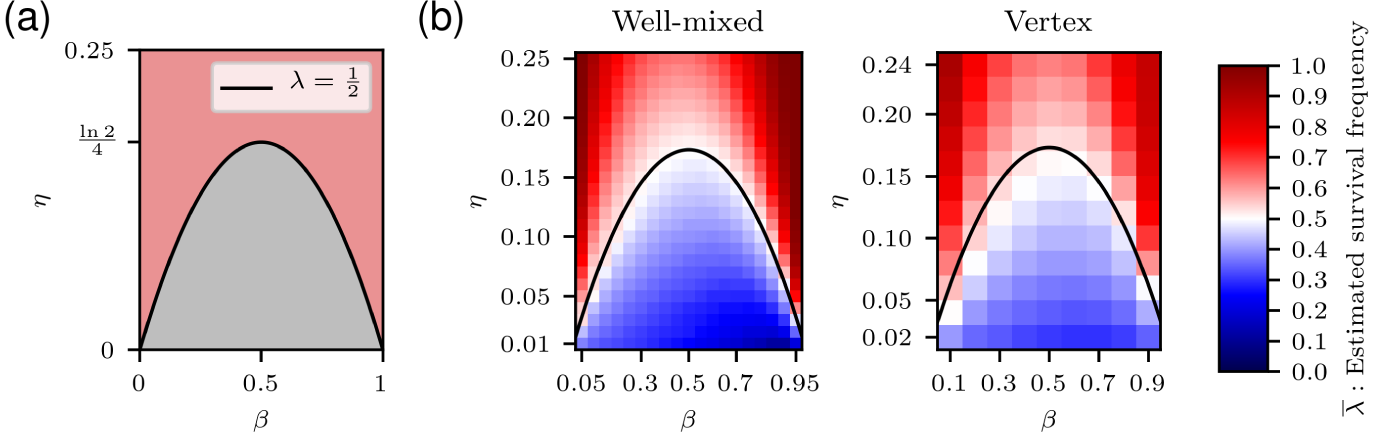
Homotypic proliferation regimes. (a) Diagram of homotypic proliferation regimes. The homotypic viability curve is given by Equation (32). Red: Viable Regime. Grey: Nonviable Regime. (b) Estimated homotypic survival frequency, 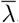, defined in Equation (33), for the well-mixed and vertex-based models. The homotypic viability curve is plotted using a black line.

#### 4.1.3. Computational validation of homotypic proliferation regimes

We use computational simulation to determine whether the viability of homotypic populations *in silico* matches the homotypic proliferation regimes as predicted by the homotypic viability curve. Further details are provided in Section S5 of the supplementary material.

For each simulation *k*, we computed the homotypic survival frequency, denoted by 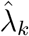, using Equation (1). For every unique parameter set, we averaged the homotypic survival frequency as

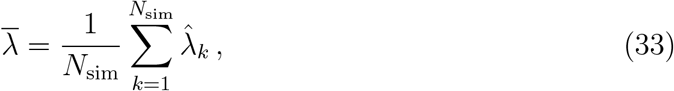

where *N*_sim_ is the number of simulations for the given parameter set.

We expect that nonviable populations tend to have a survival frequency below a half, i.e. 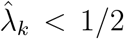, and vice versa for viable populations. Figure 4(a) predicts that cell types below the homotypic viability curve are nonviable and cell types above the curve are viable. To verify these predictions, we visualise 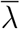 in Figure 4(b) for both the well-mixed and vertex-based models.

The left-hand plot in Figure 4(b) shows that the observed border between nonviable and viable regimes closely matches predictions for the well-mixed model. We see that for small *η* values, the survival frequency is asymmetrical with respect to *β*, with higher survival frequencies for *β <* 1*/*2 than *β >* 1*/*2. The reason for this discrepancy is discussed in Section S4.5. In short, for low *η* values, the rate of apoptosis is so high that the limiting factor is the number of cells susceptible to apoptosis, rather than the survival probability. For small *β* values, cells spend less time in G1 phase and are therefore susceptible for a shorter amount of time.

The right-hand plot in Figure 4(b) also shows good agreement between theory and simulations for the vertex-based model, although the border is less finely resolved than in the well-mixed case. We also observe the same asymmetry for small *η* values as seen in the well-mixed model.

### 4.2. Heterotypic populations

In Section 4.1, we derived an expression for the survival probability of cells in a homotypic population and used it to characterise the homotypic proliferation regimes. We take a similar approach to heterotypic populations in this section, deriving the heterotypic survival probability (Section 4.2.1) in order to map out the heterotypic proliferation regimes. However, unlike the homotypic case, the heterotypic survival probability cannot be approximated by a constant. We therefore need to define two additional quantities before we can characterise the dynamics of heterotypic populations.

Specifically, in Section 4.2.2 we define the *heterotypic survival difference*, which quantifies the difference in survival probability between competing cell types with respect to each other, and in Section 4.2.3, we define the *homotypic survival difference*, which quantifies the difference in survival probability of cells in heterotypic conditions with respect to homotypic conditions. We make use of both quantities in Section 4.2.4 to classify the different types of interactions that can occur in heterotypic populations. After analysing these derived quantities, we are able to derive the heterotypic proliferation regimes in Section 4.2.5, which we validate computationally using the well-mixed and vertex-based models in Section 4.2.6. Finally, in Section 4.3, we pull together the analyses from Section 4.1 and this section to characterise the competition regimes.

Similarly to Section 2, we create a heterotypic population in the G2 death signal model by splitting the cell population into two cell types, denoted A and B. Each cell type has its own cell cycle model, Ψ_*A*_(*t*) and Ψ_*B*_(*t*), death signal function, *f*_*A*_(*t*) and *f*_*B*_(*t*), and death threshold, *T*_†,*A*_ and *T*_†,*B*_. We assume that the cell cycle models and death signal functions are identical in both cell types, except in their parameters. With Ψ(·) as the common cell cycle model, the cell cycle models are thus parameterised as Ψ_*A*_(*t*) = Ψ(*t* ; *t*_G1,*A*_), Ψ_*B*_(*t*) = Ψ(*t* ; *t*_G1,*B*_). Similarly, the death signal functions are parameterised as *f*_*A*_(*t*) = *c*_*A*_*g*(*t*), *f*_*B*_(*t*) = *c*_*B*_*g*(*t*), with *g*(*t*) as defined in Equation (22).

#### 4.2.1. Heterotypic survival probability

In this section, we generalise the ergodic approximation, introduced in Section 4.1.1, to obtain expressions for the heterotypic survival probabilities of cell types A and B. We demonstrated for homotypic populations that the proportion of the cell cycle spent in G1 phase, *β*, is an important nondimensional parameter in determining the survival probability. Hence, in analogy with Equation (23), we define *β*_*A*_ and *β*_*B*_ for heterotypic populations such that:

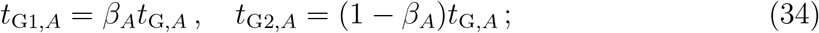

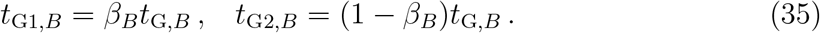

Furthermore, we assume that the ergodic property holds for both cell types separately. For cell type A, we have

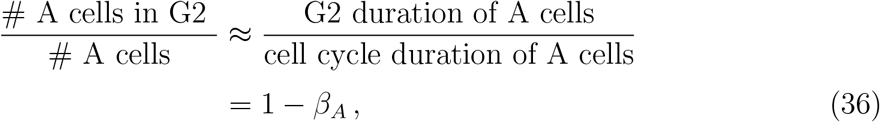

and an analogous expression can be derived for cell type B. We denote the number of A-type and B-type cells with *n*_*A*_(*t*) and *n*_*B*_(*t*), respectively, so that we can write the fraction of cells in G2 phase for the whole population as

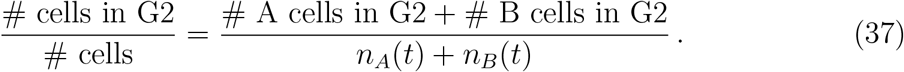

We substitute Equation (36) and its analogue for cell type B to obtain

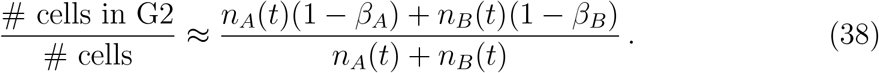

To simplify notation, we define the weighted average

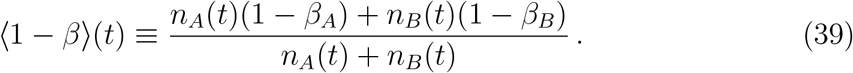

Assuming that the population is well-mixed, i.e. that Equation (25) holds, we can approximate *g*(*t*) as

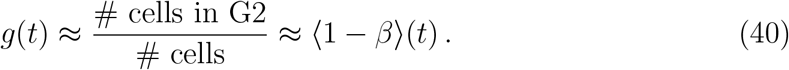

For cell type A, the death signal is thus approximated as *f*_*A*_(*t*) = *c*_*A*_*g*(*t*) ≈ *c*_*A*_ ⟨ 1 − *β* ⟩; (*t*). Note that the quantity ⟨1−*β*⟩ (*t*) is not constant with respect to time because it depends on *n*_*A*_(*t*) and *n*_*B*_(*t*). This is unlike the homotypic case (Section 4.1.1), where the death signal is approximated by the constant quantity 1 − *β*. Therefore, even with the ergodic approximation we cannot derive an exact heterotypic survival probability. Nonetheless, we can define the *instantaneous* heterotypic survival probability at time *t* as the survival probability of a cell *assuming* a constant death signal of magnitude *f*_*A*_(*t*), i.e.

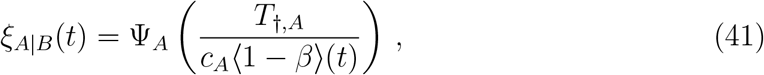

where we use the symbol *ξ*_*A*|*B*_(*t*) to denote the instantaneous survival probability at time *t* for cell type A in a heterotypic population with cell type B. Similarly, for cell type B, we have

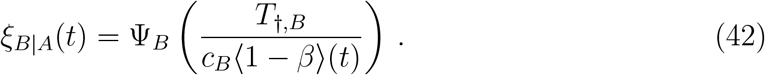

In order to derive the instantaneous heterotypic survival probability for the exponential cell cycle model in particular, we first define the dimensionless parameters

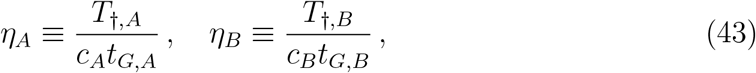

in analogy with Equation (30). We can then derive that the instantaneous heterotypic survival probabilities are

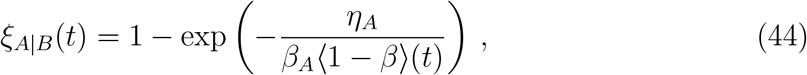

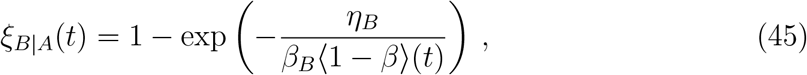

for cell types A and B, respectively. For brevity, we omit the word “instantaneous” going forward and use the symbols ⟨1 − *β*⟩ and *ξ*_*A*|*B*_ instead of ⟨1 − *β*⟩ (*t*) and *ξ*_*A*|*B*_(*t*), except when we wish to emphasise their time dependence. Furthermore, in the rest of the paper we will assume an exponential cell cycle model, unless stated otherwise.

Comparing the expressions for the heterotypic survival probability and the homotypic survival probability (Equation (31)), we see that they are almost identical, except that the weighted average ⟨1 − *β*⟩ is used instead of 1 − *β*. We note that if *n*_*B*_ = 0, then ⟨1 − *β*⟩ = 1 − *β*_*A*_ and vice versa for *n*_*A*_ = 0. In other words, when one cell type is absent, we recover the homotypic survival probability of the other cell type.

#### 4.2.2. Heterotypic survival difference

Even though the instantaneous heterotypic survival probabilities *ξ*_*A*|*B*_(*t*) and *ξ*_*B*|*A*_(*t*) change over time, in this section we show that the sign of their difference is invariant with respect to system state, and only depends on model parameters. This enables us to predict which cell type in a heterotypic population has the highest survival probability.

We define the **heterotypic survival difference** between cell types A and B as

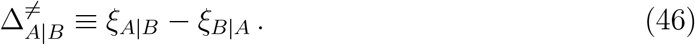

The sign of the heterotypic survival difference tells us which cell type is at a proliferative advantage. If 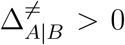, then we say that A-type cells are **winner cells** and B-type cells are **loser cells**, and vice versa for 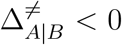 . Moreover, if 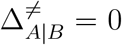, we say that the cell types are in **coexistence**, since neither cell type has a proliferative advantage over the other.

We define winners and losers here in a weak sense; if the population were to reproduce indefinitely, the winner cells would come to dominate the heterotypic population. It is not specified whether the loser population is eliminated. The classical definition of winners and losers, however, is based on the stronger condition of loser elimination. In Section 4.3, we will refine our terminology and differentiate winners and losers into more precise categories, which include classical winners and losers.

We also note that this definition of winners and losers relies on the assumption that *t*_G,*A*_ = *t*_G,*B*_, such that differences in survival probability alone determine relative proliferative success. In the general case, however, differences in the total cell cycle duration can also affect the dynamics of heterotypic populations. For instance, a cell type with a lower survival probability may become more abundant than the competing cell type by dividing more rapidly. However, for the sake of simplicity we do not consider such cases in this paper, and instead characterise population dynamics solely in terms of survival probabilities.

To obtain an expression for the sign of 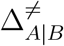, we substitute Equations (44) and (45) into Equation (46) and rearrange to give

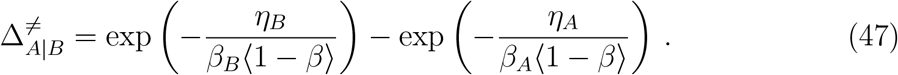

Since exp(·) is a monotonically increasing function, we have sgn(exp(*x*) − exp(*y*)) = sgn(*x* − *y*). Applying the sign function thus yields

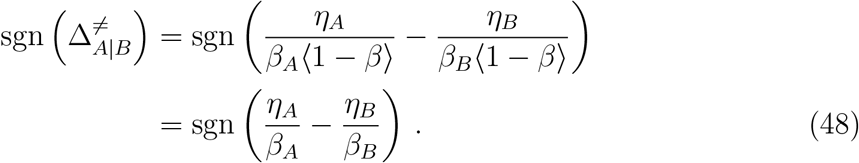

To interpret Equation (48), we note that *η* and *β* both affect a cell’s sensitivity to the death signal. Increasing *η* corresponds to a higher death threshold, and thus a lower sensitivity, and decreasing *β* shortens the time spent in G1 phase, during which a cell is vulnerable to competition-induced apoptosis. This suggests that we can interpret *η/β* as a cell’s tolerance to death signals. Therefore, Equation (48) states that the relative tolerance to death signals determines winner/loser status, with the most tolerant cell type becoming the winner.

Since the sign of 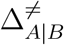 depends only on model parameters, we can partition the parameter space into two regions in which 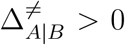 and 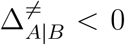, respectively. We define the **coexistence curve** for fixed *β*_*B*_ and *η*_*B*_ as the curve in (*β*_*A*_, *η*_*A*_)–space that satisfies 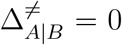 . From Equation (48), we derive that the coexistence curve is given by

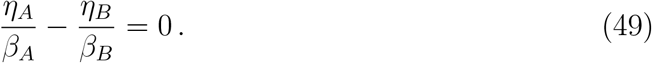

We validate this result using simulations of the well-mixed and vertex-based models in Section S6 of the supplementary material.

#### 4.2.3. Homotypic survival difference

The heterotypic survival difference does not indicate that a competitive interaction, or indeed any interaction, is taking place. After all, co-culturing two cell types that do not interact at all but have different intrinsic survival probabilities would result in a nonzero heterotypic survival difference. In this section, however, we describe a metric that quantifies changes in survival probability resulting from heterotypic interactions. In particular, we define the **homotypic survival difference** as

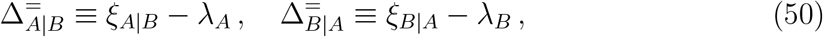

for cell types A and B, respectively. The homotypic survival difference compares the fitness of a cell type in a heterotypic environment to its fitness in a homotypic environment.

The sign of the homotypic survival difference indicates whether a cell type is more or less fit as a result of the heterotypic interaction, compared to homotypic conditions. If 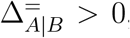, then we say that cell type A is more fit when competing with cell type B, and vice versa for 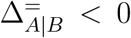. A positive homotypic survival difference indicates that the cell type benefits from the interaction. This does not mean, however, that the interaction is mutualistic, since in that case both cell types would need to benefit from the interaction (i.e. 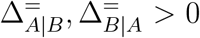. We show below that such an interaction is impossible in the G2 death signal model. Finally, if 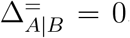, then we say that cell type A is in **neutral competition** with cell type B, since the presence of cell type B does not produce a net change in the fitness of cell type A.

Focusing our derivation on the homotypic survival difference of cell type A, we apply the sign function to give

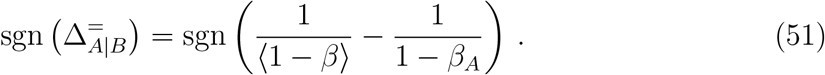

We expand ⟨1 *− β*⟩; (*t*) to give

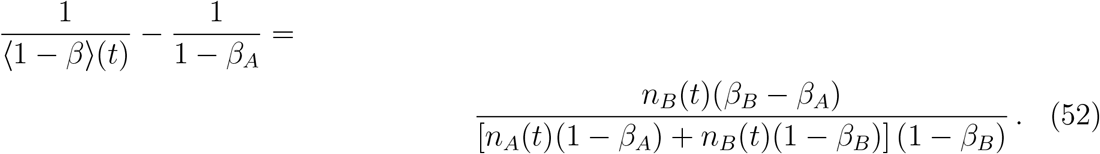

The denominator of the right-hand side is strictly positive, so we only need to consider the sign of the numerator. Equation (52) indicates that the sign of 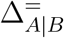 is dependent on the system state. In the degenerate case of *n*_*B*_(*t*) = 0, we are reduced to a homotypic population composed solely of A-type cells, and thus 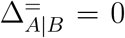. However, if we limit our scope to the heterotypic case, i.e. *n*_*A*_(*t*), *n*_*B*_(*t*) *>* 0, we can rewrite Equation (51) as

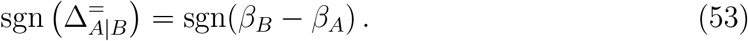

For cell type B, we derive an analogous expression:

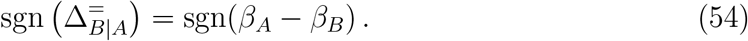

Comparing Equations (53) and (54), we derive the following identity:

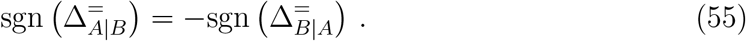

In other words, the homotypic survival differences of two competing cell types have opposite signs. Hence, one cell type’s loss is another cell type’s gain, and a mutualistic relationship is impossible.

For the heterotypic survival difference (Section 4.2.2), we factored out the death signal, ⟨1*−β*⟩;, to find an expression for the sign of 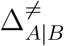 and found that winner/loser status is determined by the difference in *tolerance* to death signals. Here, in contrast, we factored out the tolerance to death signals, *η/β*, to find that the sign of the homotypic survival difference depends on the difference in *β*. Under the ergodic approximation (Sections 4.1.1 and 4.2.1), a larger value of 1 *− β* corresponds to a greater death signal. This is because the amount of time spent in G2 phase, during which cells emit death signals, is proportional to 1 *− β*. This suggests that we can interpret 1 *− β* as the cell’s emission rate of death signals. Rewriting Equation (53) as

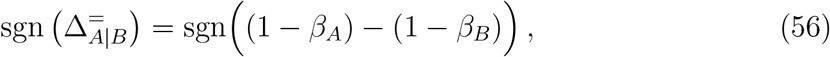

shows that the sign of the homotypic survival difference is determined by the difference in *emission* of death signals. In particular, cell type A fares better in heterotypic conditions if cell type B has a lower emission of death signal than cell type A, and vice versa.

Equations (53) and (54) show that the signs of 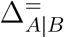 and 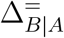 are independent of the system state, except in the degenerate homotypic cases *n*_*A*_(*t*) = 0 and *n*_*B*_(*t*) = 0. We can therefore partition the parameter space into two regions: one where 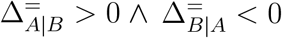, and one where 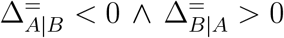. We define the **neutral competition curve** as the curve in (*β*_*A*_, *η*_*A*_)–space that satisfies 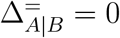 for fixed values of *β*_*B*_ and *η*_*B*_. From Equation (53), we derive that the neutral competition curve is given by

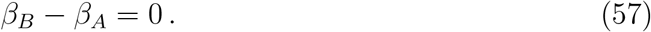

We validate this result using simulations of the well-mixed and vertex-based models in Section S7 of the supplementary material.

#### 4.2.4. Classification of competitive interactions

In Sections 4.2.2 and 4.2.3, we defined the heterotypic and homotypic survival differences, respectively. The former relates the difference in survival probability between competing cell types in heterotypic conditions, while the latter relates the difference compared to homotypic conditions. In this section, we construct a classification of competitive interactions based on these quantities.

Enumerating the signs of the homotypic and heterotypic survival differences, combined with the identity 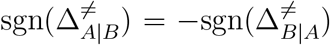 (see Equation (55)), we obtain nine types of competitive interactions (Table 4). After accounting for the fact that cell type labels are arbitrary, we can group these types into five distinct categories:

**Table 4:** Classification of competitive interactions based on the heterotypic survival difference, 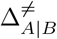defined in Section 4.2.2, and the homotypic survival difference, 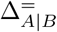, defined in Section 4.2.3.

| $\Delta_{A\vert B}^{\neq} \backslash \Delta_{A\vert B}^{\equiv}$ | + | 0 | − |
| --- | --- | --- | --- |
| + | A direct winner<br>B direct loser | A neutral winner<br>B neutral loser | A indirect winner<br>B indirect loser |
| 0 | Coexistence | Neutral coexistence | Coexistence |
| − | A indirect loser<br>B indirect winner | A neutral loser<br>B neutral winner | A direct loser<br>B direct winner |

##### Neutral coexistence 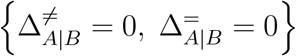

This is the degenerate case where neither cell type has a relative survival advantage, and both cell types have the same survival probability as in homotypic conditions. The competitive interaction is neutral because there is no effect on either cell type’s absolute fitness, and the cell types coexist because they have the same fitness.

##### Coexistence 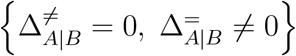

The cells experience a change in absolute fitness compared to the homotypic environment, but there is no relative survival advantage for either cell type. Therefore, neither cell type dominates.

##### Neutral competition 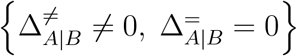

The nonzero heterotypic survival difference means that there is a difference in relative fitness. Thus, winners and losers emerge, with the winner cell type dominating the population. However, neither cell type experiences a difference in absolute fitness compared to homotypic conditions.

##### Indirect competition 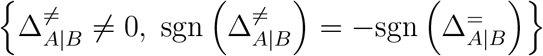

As in neutral com-petition, winners and losers emerge from the competitive interaction. The sign of the homotypic survival difference is nonzero and opposite to the sign of the heterotypic survival difference, which means that the losers experience an increase in absolute fitness compared to homotypic conditions, and the winners experience a decrease.

##### Direct competition 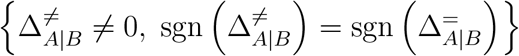

Similar to the other types of competition, the population splits into winner and loser cells. In contrast to indirect competition, however, the homotypic survival difference has the same sign as the heterotypic survival difference, meaning that the winners are fitter than in the homotypic environment, and the losers less fit.

All types of competition involve one cell type (the winners) becoming more abundant than the other cell type (the losers). The distinction between types is based on the change in fitness experienced by the winners and losers compared to homotypic conditions. In neutral competition, there is no change in fitness for either the winners or losers. In indirect competition, the winners become less fit and the losers more fit, potentially leading to a scenario where a previously nonviable loser cell type is “rescued” by the interaction with the winner cell type and becomes viable. In direct competition, the winners become more fit and the losers less fit, potentially leading to a previously viable loser cell type becoming nonviable as a result of the interaction, which is one of the cell competition criteria. We therefore expect any competitive outcomes to be the result of direct competition.

As discussed previously, we can partition cross sections of parameter space using the coexistence curve and the neutral competition curve. In Figure 5, we plot these curves in (*β*_*A*_, *η*_*A*_)–space for fixed values of *β*_*B*_ and *η*_*B*_. The curves translate to straight lines, on which we find the coexistence and neutral competition regimes. Furthermore, we find the **neutral coexistence point** at their intersection, i.e. *β*_*A*_ = *β*_*B*_ and *η*_*A*_ = *η*_*B*_, which corresponds to the degenerate case where the competing cell types have identical parameters. Finally, we see that the curves divide the cross section into four sectors, with the top left and bottom right sectors corresponding to direct competition, and the top right and bottom left sectors corresponding to indirect competition.

**Figure 5.**
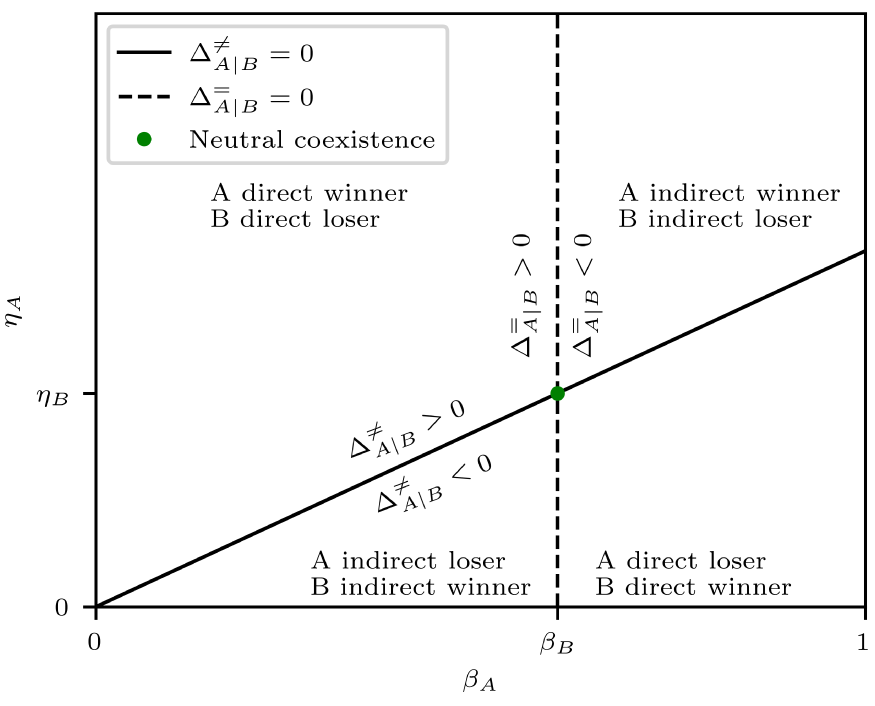
Diagram situating the different types of competitive interactions in (*β*_*A*_, *η*_*A*_)–space, given fixed values for *β*_*B*_ and *η*_*B*_. The full and dashed lines correspond to the coexistence and neutral competition curves, respectively. The green dot corresponds to the neutral coexistence point.

#### 4.2.5. Heterotypic proliferation regimes

While introducing the heterotypic survival difference in Section 4.2.2, we defined winners and losers in a weak sense based on which cell type is more prolific. Although this is an important precondition for cell competition, the cell competition criteria, as defined in Section 1.1, are based on the viability of the competing cell types, not their relative abundance. Thus, in this section we investigate the viability of winners and losers, ultimately deriving the heterotypic proliferation regimes. In Section 4.3, we use these results to arrive at a more comprehensive definition of winners and losers.

Regardless of the type of competitive interaction, winners (in the proliferative sense) become the dominant species in the population over time by definition. Therefore, we expect that the population-weighted average death signal, ⟨1 *− β*⟩; (*t*), approaches the intrinsic death signal of the winning cell type. Assuming for now that cell type A is the winner, i.e. 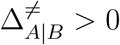, we have

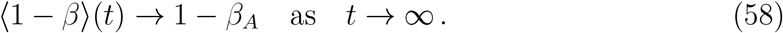

Hence, when considering the long-term behaviour of the population, we can substitute 1 *− β*_*A*_ for (1 *− β)* into the heterotypic survival probability for cell types A and B to obtain the **asymptotic survival probabilities**:

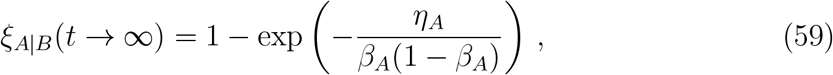

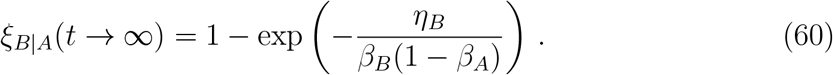

Comparing Equation (59) with Equation (31), we find that the asymptotic survival probability of cell type A is equal to its homotypic survival probability, *λ*_*A*_. The heterotypic viability of winners is thus determined by their homotypic viability. We denote the right-hand side of Equation (60) as

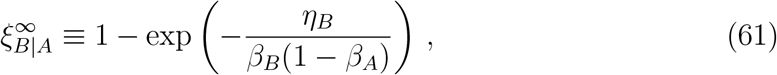

so that we can write the asymptotic survival probabilities more succinctly as

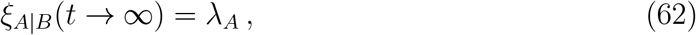

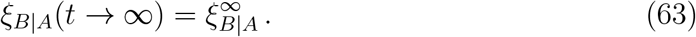

Conversely, if cell type B is the winner, i.e. 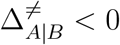, we have

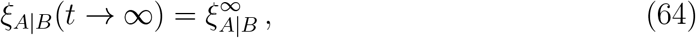

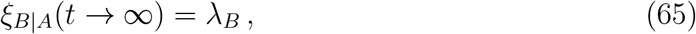

where 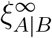 is defined analogously to Equation (61).

We can now use the asymptotic survival probability to characterise the viability of competing cell types in a heterotypic population. Assuming that cell type A is the winner, we distinguish between the following outcomes:

**Case** 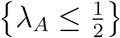

If the winner cells are not viable, then the losers are also not viable, since they have, by definition, a lower survival probability than the winners. Thus, both winners and losers go extinct.

**Case** 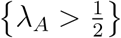

The winner cells are homotypically viable and therefore remain viable. Whether or not the losers are viable depends on 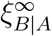 .

**Subcase** 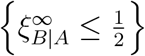

The loser cells are heterotypically nonviable and are eliminated from the tissue.

**Subcase** 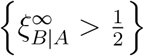

The losers are heterotypically viable and persist in the tissue.

We thus have three distinct proliferation regimes for 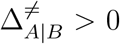. Three analogous proliferation regimes exist for 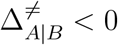, for a total of six proliferation regimes overall. We cannot visualise four-dimensional (*β*_*A*_, *η*_*A*_, *β*_*B*_, *η*_*B*_)–space directly, so we first provide an outline of the proliferation regimes, and then sketch them in cross sections for particular values of *β*_*B*_ and *η*_*B*_.

Firstly, the **coexistence hypersurface** 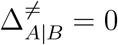 divides the parameter space into two subspaces, 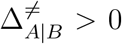 and 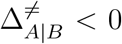, where cell types A and B are the respective winners. Secondly, for 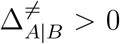, we have two regions where *λ*_*A*_ *>* 1*/*2 and *λ*_*A*_ *<* 1*/*2, respectively. The boundary is given by the **A winner viability hypersurface** *λ*_*A*_ = 1*/*2. The region in which the winner is viable, i.e. *λ*_*A*_ *>* 1*/*2, is further split into two parts, based on whether the loser is viable 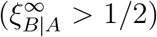 or nonviable 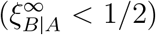, by the **B loser viability hypersurface** 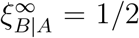. We divide the subspace 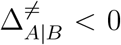, where cell type B is the winner, in an analogous manner. Hence, in total there are five hypersurfaces that delineate the heterotypic proliferation regimes: the coexistence hypersurface, two winner viability hypersurfaces and two loser viability hypersurfaces.

We visualise the heterotypic proliferation regimes using cross sections for particular values of *β*_*B*_ and *η*_*B*_ in (*β*_*A*_, *η*_*A*_)–space. In these cross sections, the hypersurfaces become the following curves:

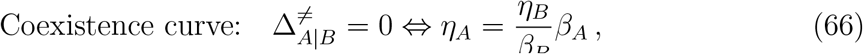

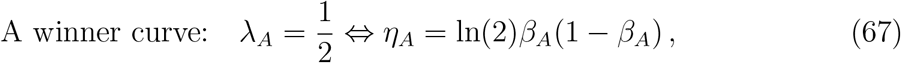

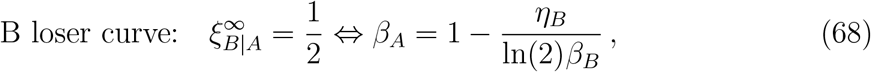

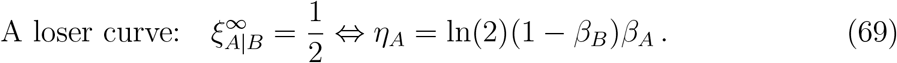

The B winner viability hypersurface does not map onto a curve in (*β*_*A*_, *η*_*A*_)–space because it depends only on *β*_*B*_ and *η*_*B*_. We therefore consider the cases *λ*_*B*_ *<* 1*/*2 and *λ*_*B*_ *>* 1*/*2 in separate cross sections.

If *η*_*B*_*/β*_*B*_ *>* ln(2), then Equation (68) does not have a solution for positive *β*_*A*_, hence the B loser viability curve does not appear in cross sections for which this is the case. We therefore consider this case in a separate cross section. It can be easily verified that *η*_*B*_*/β*_*B*_ *>* ln(2) implies *λ*_*B*_ *>* 1*/*2, so we only need to consider three distinct cross sections (Figure 6(a)):

**Figure 6.**
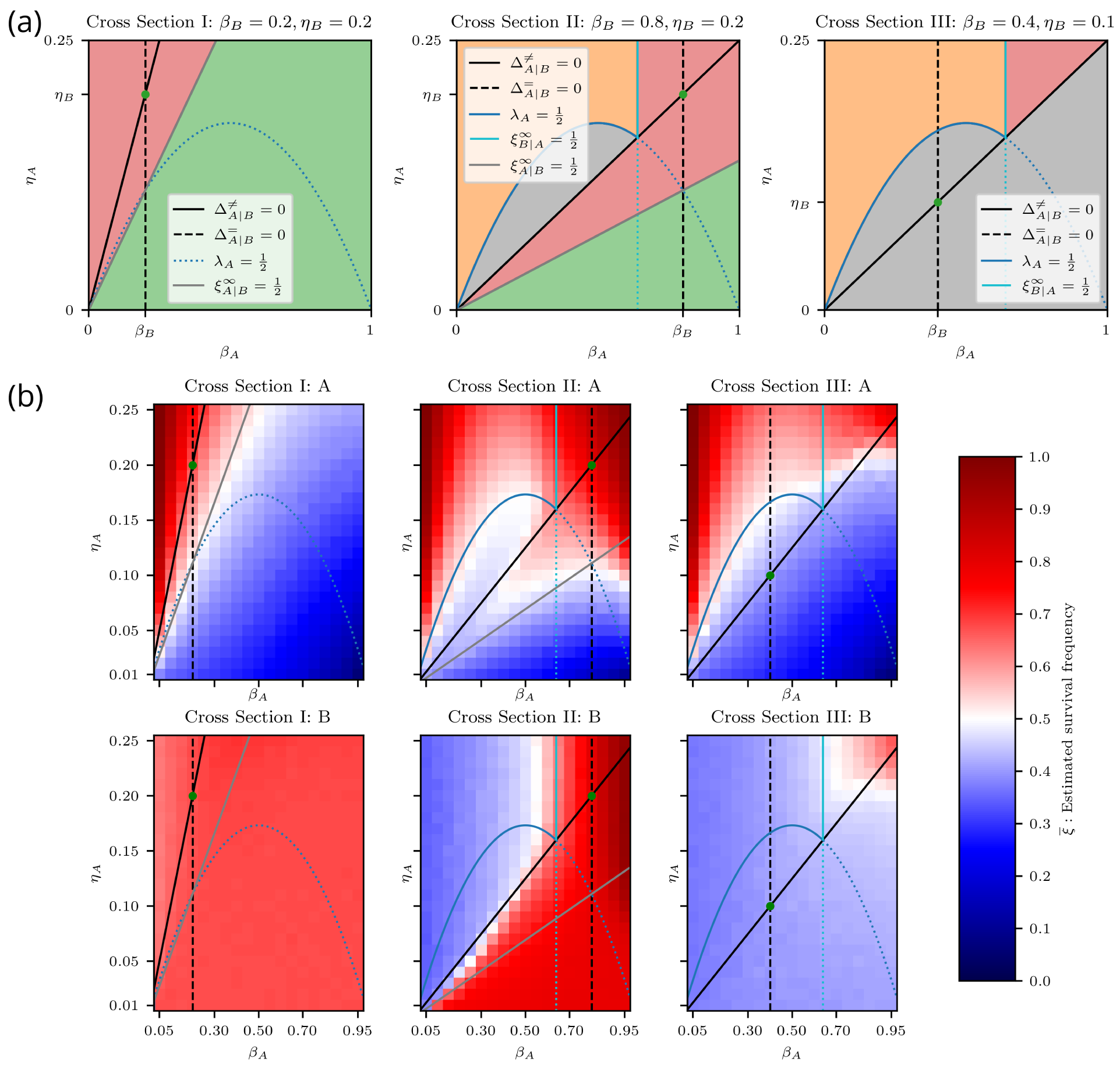
Heterotypic proliferation regimes: diagram and well-mixed results. (a) Diagrams for Cross Sections I, II, and III, situating the different heterotypic proliferation regimes. The green dot corresponds to the neutral coexistence point. Grey: cell types A and B are nonviable. Green: cell type A is nonviable, cell type B is viable. Orange: cell type A is viable, cell type B is nonviable. Red: cell types A and B are viable. (b) Estimated heterotypic survival frequency of cell types A and B using the well-mixed model. The top row displays the estimated heterotypic survival frequency of cell type A, 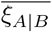, defined in Equation (70). The bottom row displays the estimated heterotypic survival frequency of cell type B, 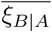, also defined in Equation (70). All curves are the same as in (a).

*Cross Section I {β*_*B*_ = 0.2, *η*_*B*_ = 0.2*}*. This cross section satisfies *η*_*B*_*/β*_*B*_ *>* ln(2). We see three distinct regimes. Above the coexistence curve, both cell types are viable, with cell type A as the winner. Between the coexistence curve and the A loser viability curve, cell type B is the winner and both cell types are viable. Below the A loser viability curve, only cell type B is viable. We note that there are no values of *β*_*A*_ and *η*_*A*_ for which cell type B is nonviable. Therefore, regardless of the competing cell type, cell type B is always viable.

*Cross Section II {β*_*B*_ = 0.8, *η*_*B*_ = 0.2*}*. This cross section satisfies *η*_*B*_*/β*_*B*_ *<* ln(2) and *λ*_*B*_ *>* 1*/*2. We identify five distinct regimes. Below the coexistence curve, we see the same two regimes as in Cross Section I. The wedge-shaped region between the coexistence curve and the A loser viability curve is particularly interesting because it partly overlaps with the area under the homotypic viability curve of cell type A. The A-type cells in this region are nonviable under homotypic conditions, but are viable when interacting with cell type B and are therefore “rescued” by the competitive interaction. This is also present in Cross Section I, but it is more visible here. We note that this region is contained within the indirect competition sector because only an indirect competitive interaction can increase the fitness of loser cells.

We see three regimes above the coexistence curve. Below the A winner viability curve, the winning A-type cells are nonviable, which renders both cell types nonviable. Above this curve, the winner A-type cells are viable. In this subspace, the survival of cell type B depends on *β*_*A*_. To the left of the B loser viability curve, the death signal emitted by cell type A is sufficiently high to eliminate cell type B, whereas, on the other side, the death signal is too weak to eliminate cell type B, so cell type B survives.

*Cross Section III {β*_*B*_ = 0.4, *η*_*B*_ = 0.1*}*. This cross section satisfies *η*_*B*_*/β*_*B*_ *<* ln(2) and *λ*_*B*_ *<* 1*/*2. Below the coexistence curve, where cell type B is the winner, both cell types are nonviable because cell type B is homotypically nonviable. Above the coexistence curve, we find the same regimes as in Cross Section II. Since cell type B is homotypically nonviable in this cross section, we note that the top right triangular region, where cell type B is heterotypically viable, corresponds to nonviable loser rescue, and thus is analogous to the wedge-shaped area discussed in Cross Section II. Similarly, this area is fully contained within the indirect competition sector.

#### 4.2.6. Computational validation of heterotypic proliferation regimes

In this section, we validate the predicted heterotypic proliferation regimes of Section 4.2.5 by conducting simulations of the well-mixed and vertex-based models. For the vertex-based model, we conducted simulations with both segregated and random initial conditions. Further details are provided in Section S8 of the supplementary material.

To estimate the survival frequency for a particular parameter set, we averaged the heterotypic survival frequencies across repeated simulations as

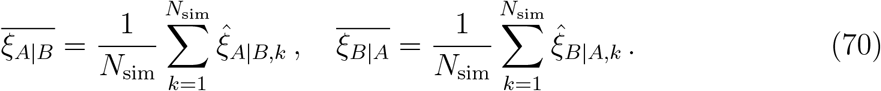

The results for the well-mixed model are given in Figure 6(b). The top and bottom rows show the survival frequency for cell types A and B, respectively. When comparing the results to Figure 6(a), we see an excellent agreement between the simulations and predictions.

The results for the vertex-based model with random and segregated initial conditions are provided in Figures 7(a) and 7(b), respectively. In Figure 7(a), we can see similar proliferation regimes as in the well-mixed case, except that the contours do not align perfectly with the predicted curves. In Cross Section II, for cell type A, we expect to see a sequence of red–blue–red–blue regions from top left to bottom right, but instead we see a gradual transition from red to blue. In addition, for high *β*_*A*_, we see red regions for cell type A that extend below their predicted limits in all cross sections.

**Figure 7.**
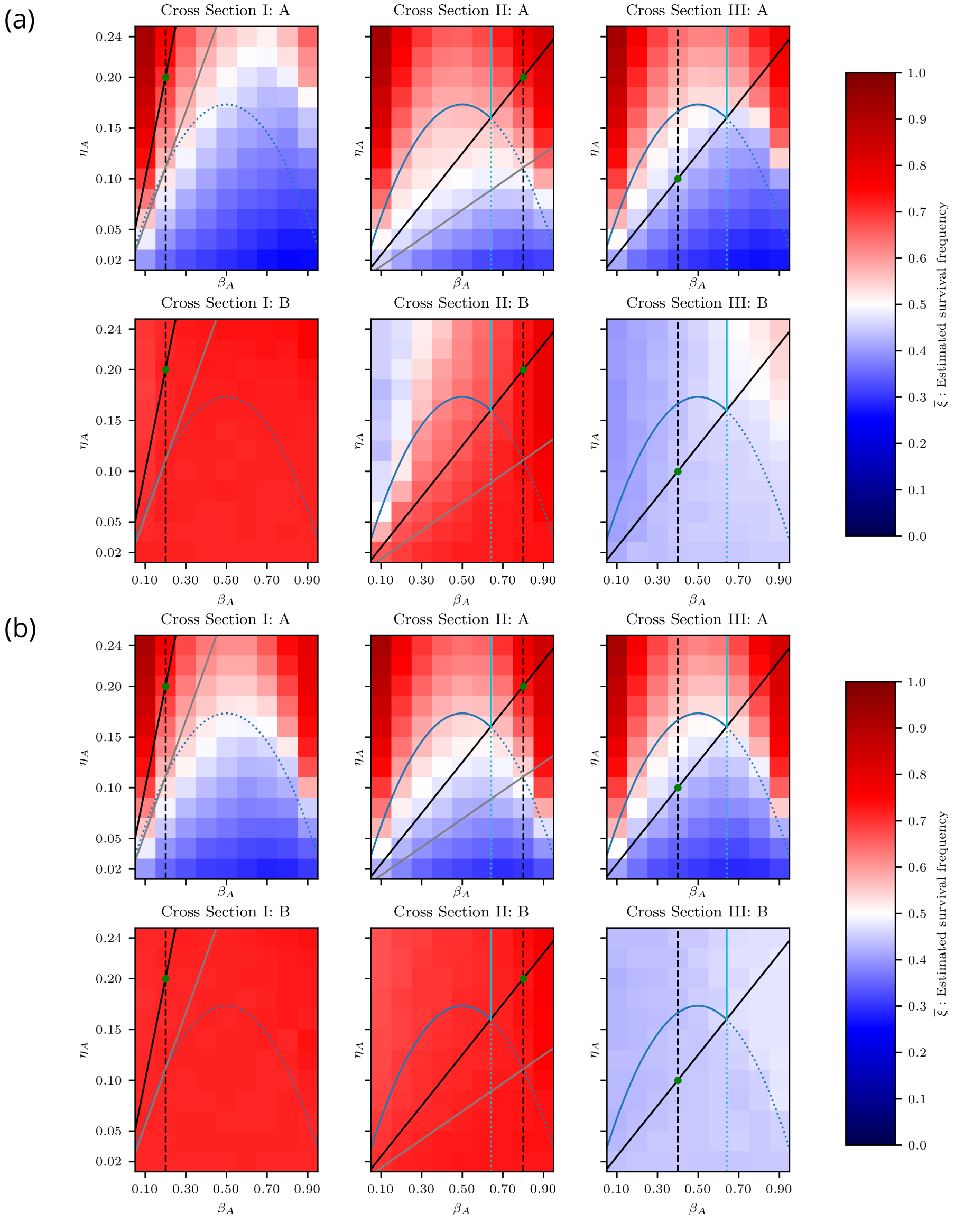
Heterotypic proliferation regimes: vertex-based results. (a–b) Estimated heterotypic survival frequency of cell types A and B using the vertex-based model with random and segregated initial conditions. See Figure 6(b) for legend. (a) Random initial conditions. (b) Segregated initial conditions.

In Figure 7(b), we see significant deviations from the predicted proliferation regimes. When comparing the plots for cell type A with the results for the homotypic proliferation regimes in Figure 4(b), we see that A-type cells essentially behave as if they were in a homotypic environment. Similarly, the heterotypic viability of cell type B matches its viability in homotypic conditions, regardless of the parameters of cell type A. These results suggest that segregated cell types behave like homotypic populations.

### 4.3. Classification of competition regimes

So far, we have systematically characterised the proliferation regimes of homotypic populations (Section 4.1.2) and heterotypic populations (Section 4.2.5). In addition, we have described and classified the different types of competitive interactions in heterotypic populations (Section 4.2.4). In this section, we integrate all these classifications into the competition regimes of the G2 death signal model, allowing us to not only apply the cell competition criteria, but also to refine and expand the known cell competition regimes.

The first condition of the cell competition criteria is that both cell types are homotypically viable, i.e. *λ*_*A*_, *λ*_*B*_ *>* 1*/*2. In order to satisfy *λ*_*A*_ *>* 1*/*2, we only consider the parameter space above the homotypic viability curve, as shown in Figure 8. To sat-isfy the viability condition for cell type B, we only consider cross sections that satisfy *λ*_*B*_ *>* 1*/*2. In particular, Cross Section III does not satisfy this condition, so we only plot Cross Sections I and II in Figure 8. We define the **homotypic viability regime** as

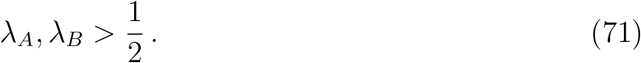

The second condition is that only one cell type remains viable when the two cell types compete. This implies a nonzero heterotypic survival difference, i.e. 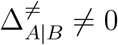, splitting the homotypic viability regime into the **coexistence regime**

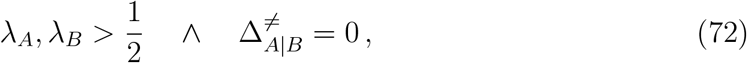

and the **competition regime**

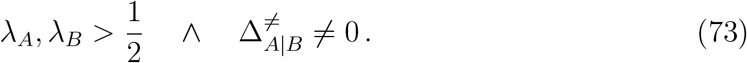

The competition regime is further subdivided according to which cell type is the winner. The G2 death signal model is symmetric with respect to swapping cell type labels, so the choice of winner or loser is arbitrary. Therefore, for ease of notation, we henceforth label the winner cell type with W and the loser cell type with L, such that 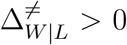 by construction.

**Figure 8.**
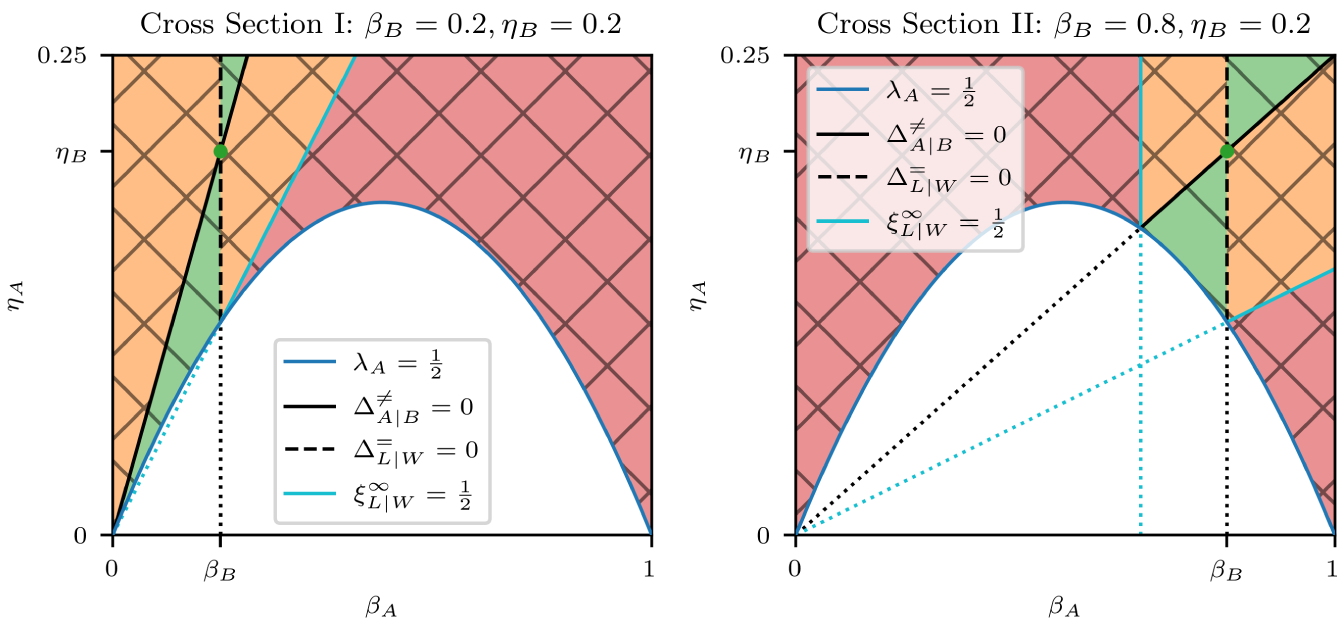
Diagrams of competition regimes for Cross Sections I and II. The green dot corresponds to the neutral coexistence point. The labels W and L are used to refer to the winner and loser cell types, respectively, i.e. W = A, L = B for 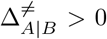 and W = B, L = A for 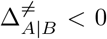. The symbol 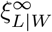 refers to the asymptotic survival probability of the loser cell type. Linear hatch: homotypic viability. Cross hatch: cell competition. Red: complete cell competition. Orange: incomplete cell competition. Green: indirect cell competition. See also Table 5 for the legend.

**Table 5:**
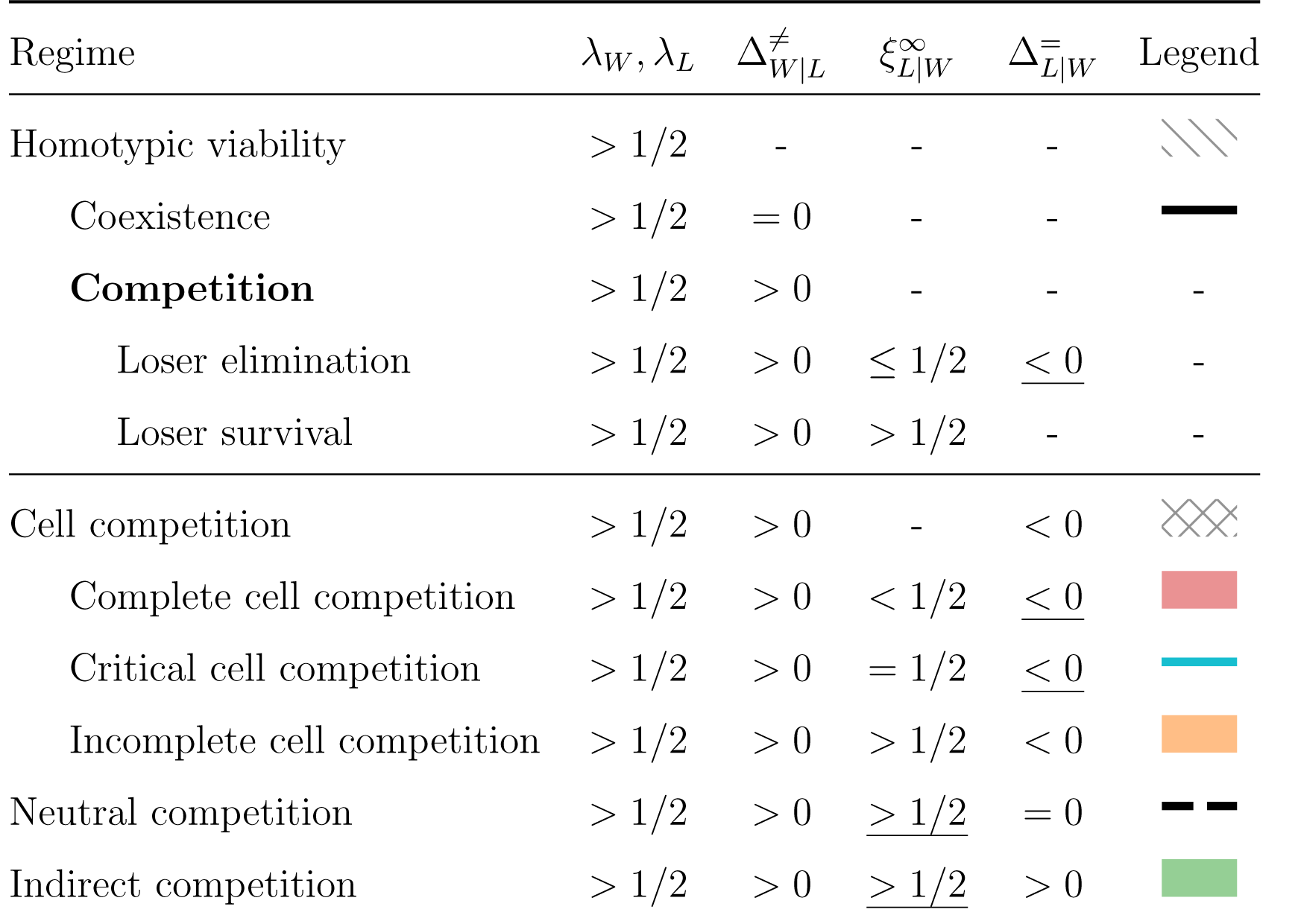
Classification of competition regimes. The competition regime (bolded) can be subdivided in two ways: loser elimination and loser survival regimes (top section), or cell competition, neutral competition, and indirect competition regimes (bottom section). The underlined conditions are implied by the other conditions on the same row. The legend column maps the regimes onto areas and curves plotted in Figure 8.

| Regime | $\lambda_W, \lambda_L$ | $\Delta_{W\vert L}^\neq$ | $\xi_{L\vert W}^\infty$ | $\Delta_{L\vert W}^\equiv$ | Legend |
| --- | --- | --- | --- | --- | --- |
| Homotypic viability | $> 1/2$ | - | - | - | |
| Coexistence | $> 1/2$ | $= 0$ | - | - | |
| **Competition** | $> 1/2$ | $> 0$ | - | - | - |
| Loser elimination | $> 1/2$ | $> 0$ | $\leq 1/2$ | <u>$< 0$</u> | - |
| Loser survival | $> 1/2$ | $> 0$ | $> 1/2$ | - | - |
| Cell competition | $> 1/2$ | $> 0$ | - | $< 0$ | |
| Complete cell competition | $> 1/2$ | $> 0$ | $< 1/2$ | <u>$< 0$</u> | |
| Critical cell competition | $> 1/2$ | $> 0$ | $= 1/2$ | <u>$< 0$</u> | |
| Incomplete cell competition | $> 1/2$ | $> 0$ | $> 1/2$ | $< 0$ | |
| Neutral competition | $> 1/2$ | $> 0$ | <u>$> 1/2$</u> | $= 0$ | |
| Indirect competition | $> 1/2$ | $> 0$ | <u>$> 1/2$</u> | $> 0$ | |

As we saw in Section 4.2.5, the viability of the winner cell type is determined by its homotypic viability, which is guaranteed by Equation (71). Therefore, we only need to impose further that the loser cell type is heterotypically nonviable, i.e. 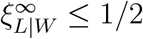. We define the **loser elimination regime** as

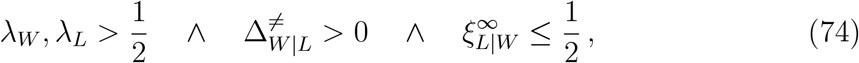

and the **loser survival regime** as

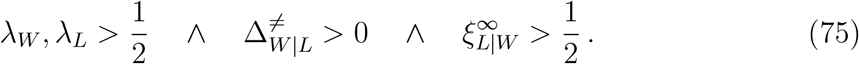

The loser elimination regime satisfies the cell competition criteria and is non-empty for the G2 death signal model. In addition, we have validated the predicted proliferation regimes with computational simulations. We therefore conclude that the G2 death signal model is capable of producing competitive outcomes.

We can further refine the competition regimes by considering, in addition, the type of competitive interaction. Figure 8 shows that the neutral competition curve, defined by 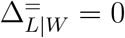, runs through the loser survival regime. We define the **neutral competition regime** as

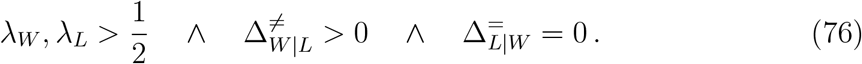

The neutral competition curve separates the loser survival regime into two subregimes where 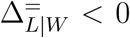 and 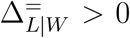, respectively. In the case of 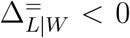, the fitness of losers is reduced by the winners, but not enough to cause loser elimination. We define this as the **incomplete cell competition regime**

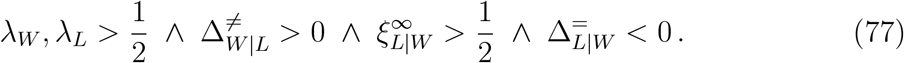

In addition, we can partition the loser elimination regime into the **complete cell competition regime**

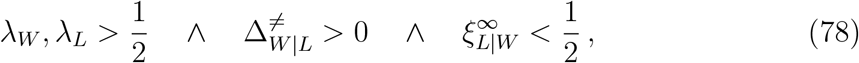

and the **critical cell competition regime**

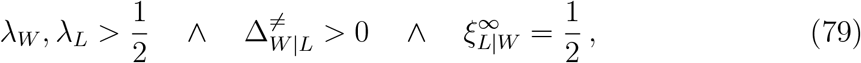

which is the threshold regime between complete and incomplete cell competition. The common feature of complete, critical, and incomplete cell competition is that the winners negatively impact the losers. We group these regimes under the **cell competition regime**

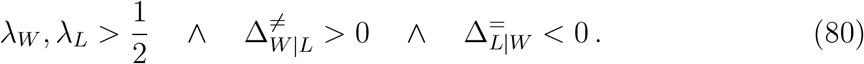

Finally, on the other side of the neutral competition curve we have 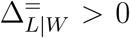, where loser cells have a higher fitness than in homotypic conditions. We denote this as the **indirect competition regime**

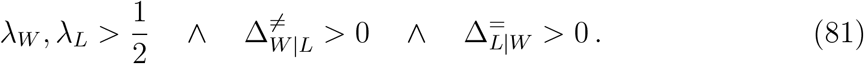

We plot the competition regimes in Figure 8 for Cross Sections I and II, and summarise them in Table 5.

The competition regimes let us discriminate between different types of winners and losers. We define *complete winners, critical winners, incomplete winners, neutral winners*, and *indirect winners* as the winner cell types in the respective competition regimes, and define different types of losers analogously. In this terminology, complete and critical winners and losers correspond to the classical definition of winners and losers in the cell competition literature.

## 5. Discussion

We stated in the introduction (Section 1) that there are two important advantages in treating winner/loser status as an emergent property rather than hardcoded identities: (i) we can test whether a given cell-based model is capable of producing competitive outcomes; and (ii), if so, analyse the conditions that give rise to competitive outcomes in that model. We demonstrated the first capability in Section 2 by showing that differences in mechanical properties alone (i.e. without a mechanism for active cell death) are insufficient to robustly generate competitive outcomes in a vertex-based model of an epithelial tissue, which agrees with experimental observations that cell competition depends on the initiation of cell death in loser cells [28].

This negative result motivated our decision to propose a modelling framework for cell competition with an active mechanism of cell death that is triggered by the exchange of death signals (Section 3). In Section 4, we introduced the G2 death signal model, in which cells only emit death signals in the G2 phase. We systematically investigated its behaviour for homotypic (Section 4.1) and heterotypic populations (Section 4.2), studying their proliferation regimes through a combination of (i) theoretical analysis based on the survival probability and (ii) computational simulation using the well-mixed and vertex-based models, ultimately culminating in the characterisation of the competition regimes in Section 4.3. Importantly, our analysis allows for a direct examination of the conditions and parameters that lead to competitive outcomes. In this section, we will interpret and discuss our findings, propose specific ideas for novel cell competition experiments, and outline potential future research directions.

### 5.1. Spatial mixing is required for cell competition

In Section 4.2.6, we observed that the occurrence of competitive outcomes in the vertex-based model depends on the initial spatial patterning of cell types. When the cell types are distributed randomly, we observe competitive outcomes, but when they are segregated, we do not. In fact, the behaviour of the segregated cell types is virtually identical to that of isolated homotypic populations. This result agrees with experimental observations that spatial mixing is required for cell competition [42], and has been replicated in other cell-based models of cell competition [13].

Our derivation of heterotypic proliferation regimes is based on the assumption that the population is well-mixed, which is only true *locally* at heterotypic clone boundaries in the vertex-based model, where cells sample the death signal of both cell types. Within clones, however, cells interact only with cells of the same type, so they behave more like a homotypic population. The degree of competition therefore depends on the amount of heterotypic contact between cell types, which is modulated by the level of spatial mixing.

### 5.2. Tolerance and emission

When we derived the heterotypic survival difference in Section 4.2.2, we found that the relative abundance of cell types in a tissue is determined by their tolerance to death signals (i.e. *η/β*). Furthermore, when we derived the homotypic survival difference in Section 4.2.3, we showed that the difference in death signal emission (i.e. 1*− β*) between two competing cell types determines the impact of the heterotypic interaction compared to homotypic conditions. Also, in Section 4.2.5, we demonstrated that loser elimination depends on the relationship between the tolerance of the loser and the emission of the winner. From these observations, we infer that tolerance to, and emission of, death signals are the fundamental cell properties driving cell competition in the G2 death signal model. Here, we present a transformation of parameters that explicitly describes the behaviour of the model in terms of tolerance and emission. We also show that the transformed parameters allow us to describe the competition regimes using intuitive and elegant expressions.

We define the **tolerance** and **emission** of cell type *X*, respectively denoted 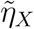 and *d*_*X*_, as follows:

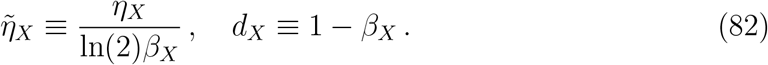

We can formulate the **homotypic viability** condition, 1*/*2 *< λ*_*X*_, using 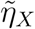 and *d*_*X*_ by substituting the homotypic survival probability (Equation (31)) and rearranging:

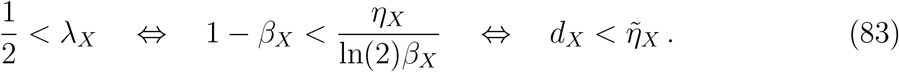

The last inequality reads as the condition that cells must have a higher tolerance than emission to be homotypically viable. The biological interpretation is that cells must be capable of tolerating the death signal that they themselves emit in order to survive as a group. The **loser elimination** condition, 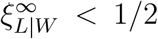, can also be expressed using tolerance and emission. Denoting the winner and loser cell types using the labels W and L, respectively, we substitute the asymptotic survival probability of the loser (Equation (61)) to obtain

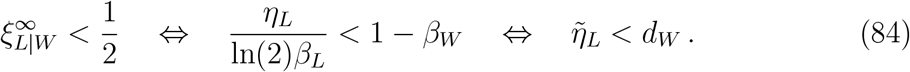

This means that winner cells must emit death signals at a rate that loser cells cannot tolerate in order to eliminate the loser cell type from the tissue.

To satisfy the cell competition criteria, we require that both cell types are homotypically viable, i.e. 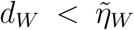 and 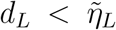, and that the loser is eliminated, i.e.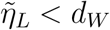. Combining these expressions, we can summarise the conditions on the model parameters such that the cell competition criteria are satisfied in a single statement:

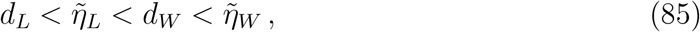

which can be read as

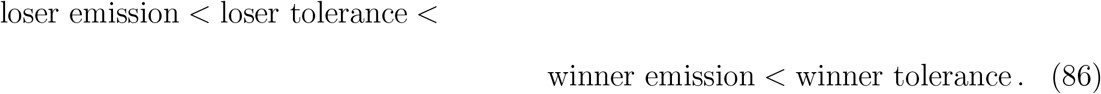

This corresponds to the **complete cell competition** regime that we defined earlier in Section 4.3. In a similar manner, we can express all the competition regimes defined in Section 4.3 in terms of tolerance and emission (compare the following with the bottom section of Table 5):

**Cell competition:** 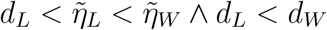.

**Complete cell competition:** 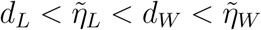.

**Critical cell competition:** 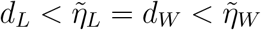.

**Incomplete cell competition:** 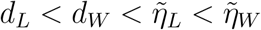 .

**Neutral competition:** 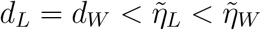 .

**Indirect competition:** 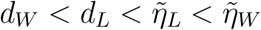 .

These relationships can be verified visually in Figure 9, which shows the competition regimes in transformed parameter space.

**Figure 9.**
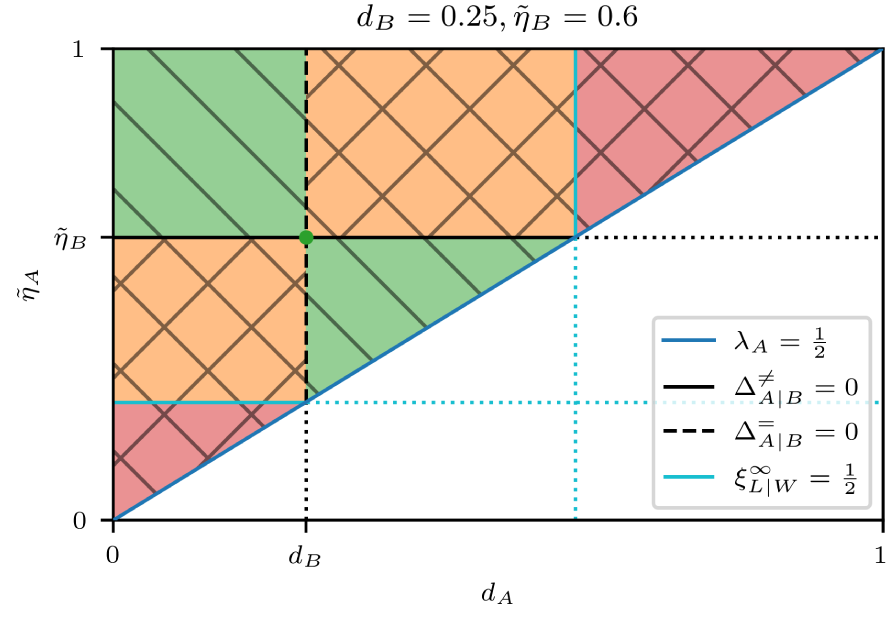
Diagram of competition regimes using the transformed parameters 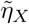 and *d*_*X*_, defined in Equation (82). The green dot corresponds to the neutral coexistence point. The same conventions apply as in Figure 8. See Table 5 for the legend.

### 5.3. The tolerance–emission model of cell competition

Based on Equation (86), we make the following biological prediction: **cell competition requires that winner cells have a higher tolerance to death signals *and* a higher rate of death signal emission than loser cells**. The implicit assumption in this statement is that cells emit and tolerate some form of death signal, which can be contact-based, ligand-based, mechanical stress-based, etc. Intuitively, regardless of the type of death signal, winners must expose losers to a sufficiently high level of death signal to eliminate them, while still being able to withstand it themselves.

Importantly, this model implies that mutations resulting in cell competition, such as *Minutes* and *Myc*, are pleiotropic because they simultaneously alter the tolerance to, and emission of, death signals. As a corollary, mutations which affect only one or neither, do *not* engender cell competition. This potentially explains why some mutations related to proliferation rates result in cell competition, and others do not [7]. In this view, the inhibition of apoptosis can be regarded as a mutation that results in an infinite tolerance, without affecting emission. Indeed, it has been shown in experiments that inhibiting apoptosis prevents cell competition [7, 28].

This observation raises the question: do mutations exist that increase the emission of death signals, without affecting tolerance? If so, they would be challenging to culture, since such mutants would not tolerate their own death signal and thus be homotypically nonviable. However, the tolerance–emission model suggests that such a mutation would be viable if it were paired with apoptosis inhibition. Our model therefore predicts that a hypothetical emission-enhancing mutation combined with apoptosis inhibition would result in a novel species of super-competitors.

#### 5.3.1. Experimental support

Experimental evidence from *Myc*-based cell competition supports the tolerance– emission hypothesis. In [43], the authors demonstrated that the ligand Spätzle is necessary for the elimination of loser cells in the *Drosophila* wing disc, forming what the authors term a “killing signal”. They also observed that Spätzle is produced in wild-type conditions at a rate that is tolerated by the wild-type cells, and that the production of Spätzle is upregulated in *Myc* super-competitors without inducing cell death in *Myc* mutants. *Myc* mutants therefore emit more death signal than wild-type cells, while simultaneously being less sensitive to it.

In this experiment, the death signal takes on the form of a diffusible death ligand. While the principles of tolerance and emission should still apply, there is an important difference with the contact-based G2 death signal discussed in Section 4; namely that death ligands can diffuse away from the site of heterotypic contact. This could potentially explain why we observe loser cell death at a distance in *Myc*-based cell competition [7], but not in *Minutes*-based cell competition. According to the tolerance–emission model, death ligand secretion is upregulated in mutant winner cells in the former case, and downregulated in mutant loser cells in the latter case.

We also find support for the tolerance–emission model in mechanical cell competition, specifically in cultures of Madine–Darby canine kidney (MDCK) cells [10]. The authors discovered that cell proliferation is in part modulated by the composition of cell types in the cellular neighbourhood. In particular, winner cells are more prolific when they are specifically surrounded by loser cells. This agrees with our observations that winner cells benefit from proximity to loser cells because loser cells emit a lower level of death signal.

#### 5.3.2. Experimental validation

To validate the tolerance–emission hypothesis, we must extrapolate the model predictions to experimental conditions that have not yet been tested. We predicted in Section 4.2.5 that homotypically nonviable loser cells can be rescued through indirect competition. This occurs when a winner cell type has a lower emission rate than the loser cell type, creating an environment in which losers can proliferate even if they are not viable on their own. The challenge in producing this outcome experimentally, however, is that we would first need to identify an intrinsically nonviable mutant cell type to assume the role of the loser. We therefore propose an alternative experiment that could potentially simulate this behaviour with known cell types.

Consider a triple co-culture where cell type A outcompetes cell type B and cell type B outcompetes cell type C. Cell types B and C are both eliminated in a background of cell type A, which mimics the intrinsic nonviability of cell types B and C. The tolerance– emission model predicts that the emission of death signals by cell type C is tolerated by cell type B. Therefore, if we inhibit apoptosis in cell type C, we expect to see: (i) C-type clones forming in an A-type background; and (ii) the survival of B-type cells exclusively inside the C-type clones. This outcome would be analogous to the rescue of a homotypically nonviable loser by indirect competition, with cell types B and C corresponding to the indirect losers and winners, respectively.

### 5.4. The function of cell competition

The prevalent hypothesis is that cell competition is a mechanism for maintaining tissue health by eliminating unfit cells. However, what is meant by “fitness” in this context is not clear [44]. The classical definition of fitness is based on *reproductive success* and early experiments indeed linked reproductive fitness to cell competition, with winner cells having higher intrinsic proliferation rates than losers [45, 46]. However, not all mutations that increase proliferation rates result in cell competition [7]. In cell competition, fitness is perhaps more accurately defined as a measure of *competitive success*, which can determined by pairwise contests between cell types. In the tolerance– emission model, competitive success is a combination of tolerance and emission, and lacks a causal relationship with proliferation rates.

Competitive fitness is therefore not the same as reproductive fitness, but then why are they often linked in practice? We speculate that differential proliferation rates are not the mechanism of cell competition, but the *target* of cell competition. Cell competition evolved to optimise reproductive fitness, but uses competitive fitness as an imperfect means to communicate it. In other words, competitive fitness serves as a proxy for reproductive fitness and evolved in a trade-off with other factors such as the costs involved in cell competition.

Furthermore, we expect that the target of cell competition depends on the function of the host tissue. In the *Drosophila* wing disc, the tissue expands from 50 to 50 000 cells in the span of four days, so the function of cell competition in this context is to optimise for reproductive fitness. MDCK cells, on the other hand, were derived from kidney tubules, so their function is to form a mechanically resilient barrier. In this case, cell competition is linked to mechanical cell compression, which we hypothesise acts as a proxy for the cell’s ability to contribute to the structural integrity of the tissue.

### 5.5. Future work

The framework presented in this paper can be applied to any cell-based model to study hypothetical mechanisms of cell competition. Moreover, the cell competition criteria are sufficiently abstract that they can potentially be translated to models of cell competition that are not cell-based, such as Lotka–Volterra models [47].

We emphasise that the death clock framework is agnostic with respect to the death signal, and that it can be used to represent different kinds of cell competition mecha-nisms. Of particular interest are diffusible ligands and mechanical compression as death signals. Studies show that cell competition in the *Drosophila* wing disc involves the use of diffusible death ligands [43, 48, 49]. A death clock model based on the secretion (i.e. emission) and recognition of death ligands is therefore an obvious next step toward a more biologically accurate representation of the cell competition process. Section 2 suggests that differences in mechanical properties alone do not robustly generate competitive outcomes in a heterotypic vertex-based model. However, they may still play a role in cell competition when paired with an active mechanism for cell death. Research indicates that mechanical compression triggers apoptosis in loser cells during mechanical cell competition [16], hence cell compression may be an appropriate death signal in this context. Further research is needed to investigate models that incorporate diffusible ligands or mechanical compression as death signals.

## 6. Acknowledgements

The authors would like to acknowledge the use of the University of Oxford Advanced Research Computing (ARC) facility [50] and GNU Parallel [51] in carrying out this work. This work was supported by the Biotechnology and Biological Sciences Research Council (BBSRC) [BB/M011224/1].

## Footnotes

1 The astute reader may note the discrepancy between the definition of viability based on survival probability versus the definition based on survival frequency (Section 1.2): *λ* = 1*/*2 is considered nonviable, whereas 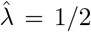 is considered viable. This subtle distinction is rooted in the theory of birth–death Markov chains but bears no significance on our argument so we will not go into it further.

