## Supplementary material for "A mathematical framework for the emergence of winners and losers in cell competition"

Thomas F. Pak, Joe M. Pitt-Francis, Ruth E. Baker

#### S1 Statistical analysis of targeted simulation suites

In Section 2.3, we ran a large parameter sweep of the mechanical model to search for competitive behaviour, and found that only 20 out of 2 809 parameter sets (less than 1%) produced positive results. Because the mechanical model is stochastic, in this section we run additional simulations to test the statistical significance of the competitive outcomes we observed. In addition, we investigate the effects of segregation on competitive behaviour. We therefore conduct two targeted simulation suites of the 20 aforementioned parameter sets: one with random initial conditions, and one with segregated initial conditions. See Figure S1(c–d) for examples of random and segregated initial conditions, respectively.

##### S1.1 Methods

As mentioned in Section 2.2, we need to run two homotypic simulations, one for each cell type, and one heterotypic simulation to sample the homotypic and heterotypic viabilities of two cell types. We call such a set of simulations a simulation triplet. In each simulation suite (random and segregated), we conducted 20 simulation triplets for each parameter set, for a total of $3 \times 20 \times 20 = 1\,200$ simulations per simulation suite. We treat each parameter set independently as a Bernoulli process. Hence, in this view, one simulation triplet corresponds to a Bernoulli trial with probability $q$ of satisfying the cell competition criteria. If a parameter set is more likely to produce a competitive outcome than not, i.e. $q > 1/2$, then we say that it gives rise to cell competition. In order to test a parameter set for competitive behaviour, we discriminate between the following hypotheses:

$$\mathbf{H_0} : q \leq \frac{1}{2}\,; \tag{S1}$$

$$\mathbf{H_1} : q > \frac{1}{2}\,. \tag{S2}$$

In other words, the null hypothesis is that the parameter set does not give rise to cell competition, i.e. $q \leq 1/2$, and the alternative hypothesis is that it does, i.e. $q > 1/2$.

We decide whether or not to reject the null hypothesis by calculating how likely it is to obtain results that are at least as extreme as the observed results under the null hypothesis. This probability is also known as the $p$-value. By "extreme", we mean results that support the alternative hypothesis rather than the null hypothesis. In this case, the more simulation triplets that match the cell competition criteria, the more extreme the observed data are. For example, observing ten matches provides stronger evidence in favour of competitive behaviour than three matches. A small $p$-value indicates that the observed results are unlikely to be obtained under the null hypothesis, supporting the alternative hypothesis, and vice versa for a large $p$-value.

#### S1.2 Results

In each simulation suite, we counted the number of competitive outcomes for each parameter set to calculate its $p$-value. In the random simulation suite, we found that six parameter sets produced a statistically significant number of positive results at a significance level of 5%. Hence, for these six parameter sets we reject the null hypothesis and conclude that they generate competitive behaviour. In contrast, only one parameter set was statistically significant in the segregated simulation suite, indicating a marked reduction in competition as a result of spatial segregation.

For visual comparison, in Figure S1 we show a pair of homotypic simulations, and heterotypic simulations in random and segregated initial conditions. We picked a parameter set that produced statistically significant competitive outcomes for random initial conditions, but not for segregated initial conditions. It is clear from the homotypic simulations that the losers are intrinsically much less prolific than the winners, and grow to a much smaller clone size over the same time period. This pattern was common to all six statistically significant parameter sets.

Comparing the heterotypic simulations, it appears that, for random initial conditions, the loser cells are extruded as a result of being surrounded by densely packed and quickly proliferating winner cells. For segregated initial conditions, however, cell type A is located at the edge of the tissue, which lets them escape the intense "growth pressure" in the interior of the tissue. Hence, from visual inspection it seems that competitive behaviour in the mechanical model is mediated by differences in homeostatic pressure, which is reminiscent of the model proposed in [1]. However, more work is needed to verify this hypothesis.

### S2 Implementation details of well-mixed model

In this section, we discuss implementation details of the well-mixed model (Section 3.4.1). We first describe the discrete operations in more depth in Section S2.1, followed by a discussion of the numerical implementation in Section S2.2.

#### S2.1 Discrete operations

The discrete division and death operations are triggered when the **division variant** and the **death variant**, respectively, are violated. The division invariant is as follows:

$$\forall \alpha = 1, \dots, N(t) : t - t_\alpha^0 < t_\alpha^* + t_{\mathrm{G2},\alpha} \,, \tag{S3}$$

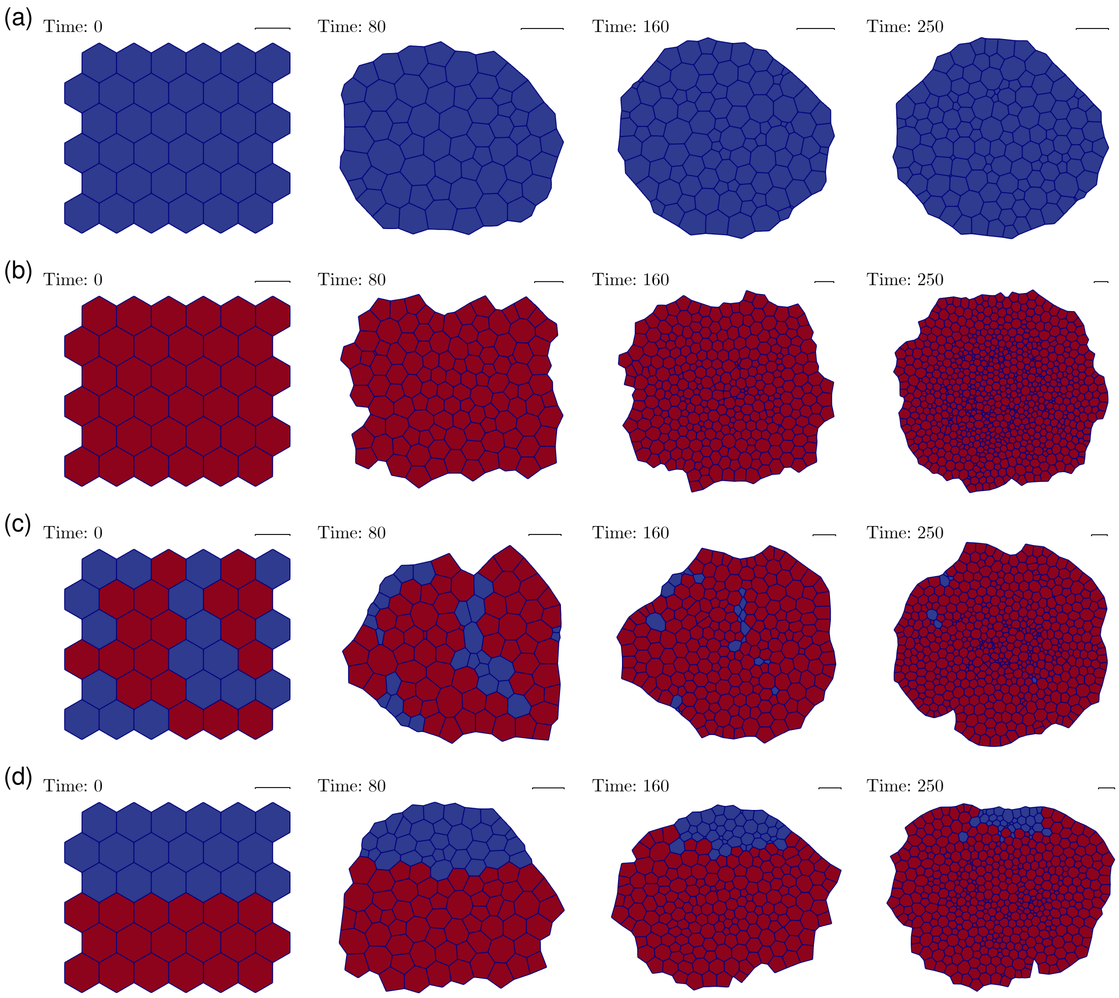


Figure S1: Example simulations of the mechanical model for parameter set: $S_A^0 = 0.981$, $K_A = 0.547$, $\Gamma_A = 0.021$, $\Lambda_{AA} = 0.097$, $t_{\mathrm{G1},A} = 26.598$, $t_{\mathrm{G2},A} = 74.522$, $S_B^0 = 1.073$, $K_B = 1.113$, $\Gamma_B = 0.055$, $\Lambda_{BB} = 0.165$, $t_{\mathrm{G1},B} = 0.560$, $t_{\mathrm{G2},B} = 48.484$, $\Lambda_{AB} = 0.145$. Cells are coloured according to their cell type. Blue: cell type A. Red: cell type B. (a–b) Homotypic simulations for cell types A and B. (a) Cell type A is homotypically viable with a final cell count of 136. (b) Cell type B is homotypically viable with a final cell count of 810. (c–d) Heterotypic simulations for random and segregated initial conditions. (c) Cell type B outcompetes cell type A with final cell counts 6 and 554 for cell types A and B, respectively. (d) Both cell types remain viable with final cell counts 36 and 471 for cell types A and B, respectively.

or, in words, "every cell's age is less than its total cell cycle duration". The death invariant is as follows:

$$\forall \alpha = 1, \ldots, N(t) : t - t_\alpha^0 < t_\alpha^* \Rightarrow \tau_\alpha(t) < T_{\dagger,\alpha}\,, \tag{S4}$$

or, in words, "for every cell in G1 phase, its death clock is below its death threshold".

These invariants must be respected at all times during the simulation, including the initial conditions. When an invariant is violated, the corresponding operation is instantly triggered. The operation is guaranteed to restore the invariant, therefore allowing the simulation to proceed while respecting the invariants.

##### S2.1.1 Division operation

As soon as the division invariant is violated, i.e. if

$$\exists\, \alpha \in 1, \ldots, N(t) : t - t_\alpha^0 = t_\alpha^* + t_{\mathrm{G2},\alpha}\,, \tag{S5}$$

or, in words, "as soon as any cell's age reaches its total cell cycle duration", division is triggered. Biologically, the mother cell splits into two daughter cells. Computationally, we perform this operation by adding a daughter cell vector and reusing the mother cell vector to construct the second daughter cell in-place.

Concretely, we update the population as[1]

$$S(t) \leftarrow S(t) \cup \{\, \mathbf{y}_{N(t)+1}(t) \,\}\,, \tag{S6}$$

and set the contents of the cell vectors $\mathbf{y}_\alpha(t)$ and $\mathbf{y}_{N(t)+1}(t)$ as below. Finally, we increment $N(t)$ to reflect the increased population count.

**Inherit cell cycle and death clock components** Both daughter cells inherit the mother cell's cell cycle and death clock components. Since we repurpose the mother cell vector $\mathbf{y}_\alpha(t)$ for the first daughter cell, we simply copy the relevant fields to $\mathbf{y}_{N(t)+1}(t)$:

$$\mathcal{C}_{N+1} \leftarrow \mathcal{C}_\alpha\,, \tag{S7a}$$
$$t_{\mathrm{G2},N+1} \leftarrow t_{\mathrm{G2},\alpha}\,, \tag{S7b}$$
$$f_{N+1}(\cdot) \leftarrow f_\alpha(\cdot)\,, \tag{S7c}$$
$$T_{\dagger,N+1} \leftarrow T_{\dagger,\alpha}\,. \tag{S7d}$$

**Sample G1 duration** We sample a G1 duration for each daughter cell from their cell cycle distribution as[2]

$$t_\alpha^* \twoheadleftarrow \mathcal{C}_\alpha\,, \tag{S8a}$$
$$t_{N+1}^* \twoheadleftarrow \mathcal{C}_{N+1}\,. \tag{S8b}$$

[1]The symbol '$\leftarrow$' represents the update operator, which updates the left operand(s) with the value of the right operand.

[2]The symbol '$\twoheadleftarrow$' represents the sample-and-update operator, which updates the left operand with a value sampled from the right operand, which must be a stochastic distribution.

**Set birth time** We set the birth times of the daughter cells as

$$t_\alpha^0, t_{N+1}^0 \leftarrow t_{\text{division}} \,, \tag{S9}$$

where $t_{\text{division}}$ is the time at which the division invariant was violated.

**Reset death clock** We reset the death clocks of the daughter cells:

$$\tau_\alpha^0, \tau_{N+1}^0 \leftarrow 0 \,. \tag{S10}$$

##### S2.1.2 Death operation

As soon as the death invariant is violated, i.e. if

$$\exists\, \alpha \in 1, \ldots, N(t) : t - t_\alpha^0 < t_\alpha^* \text{ and } \tau_\alpha(t) = T_{\dagger,\alpha} \,, \tag{S11}$$

or, in words, "as soon as the death clock of a cell in G1 phase reaches the death threshold", apoptosis is triggered.

The population is updated as

$$S(t) \leftarrow S(t) \setminus \{\, \mathbf{y}_\alpha(t) \,\} \,, \tag{S12}$$

followed by a reindexing operation to reflect the decreased population count. We also decrement $N(t)$.

#### S2.2 Numerical implementation

We implemented the well-mixed model in Python using the class `WellMixedSimulator`. This class is responsible for (i) checking the validity of the initial conditions, (ii) simulating the system forward in time by applying the death clock ODE and discrete operations, and (iii) terminating the simulation. In addition, the class records the state of the system throughout the whole simulation and returns the raw simulation data upon termination. The code for the model can be found in the following GitHub repository: `https://github.com/ThomasPak/cell-competition`.

##### S2.2.1 ODE solver

The simulation proceeds by numerically solving Equation (20) for all cells until an invariant is violated, which triggers the corresponding discrete operation. By default, we use the SciPy[3] ODE solver, `solve_ivp`, to integrate Equation (20). The `events` argument to the solver lets us specify conditions on the ODE system for which the solver terminates prematurely, i.e. before the simulation end time is reached. We use this feature to exit the ODE solver when an invariant is violated so that we can perform the corresponding discrete operation outside of the solver. After restoring the invariant, numerical integration of Equation (20) is resumed. The discrete operations are implemented by methods of the `WellMixedSimulator` class. In summary, the standard mode of operation is numerical integration within the SciPy ODE solver, punctuated by division and death events implemented in the simulator class.

[3] `https://scipy.org/`

##### S2.2.2 Optimised ODE solver

Although the SciPy ODE solver is relatively efficient, we can dispense with numerical integration entirely in special cases to gain a massive performance boost. In particular, if the death signal is constant for all cells in between discrete events, then Equation (20) can be solved exactly. Hence, the time until the next event can be computed by simple arithmetic operations. For instance, if the death signal depends only on the total cell count, then the death signal only changes with division and death events, so we can apply this optimisation.

In addition to the division and death events, we added the transition from G1 phase to G2 phase as a third type of discrete event in the simulator. This event is not strictly necessary for simulating the model, but it forces the ODE solver to terminate when a cell transitions from G1 phase to G2 phase. With this extended set of events, death signals that depend on the distribution of G1 and G2 phases (such as the G2 death signal model discussed in Section 4) are constant in between discrete events, such that the optimisation can be applied in those cases as well. We enable this optimisation by passing the Boolean flag `f_is_stepwise_constant` to the simulator.

##### S2.2.3 Pseudorandom number generator

Since G1 durations are sampled from a stochastic distribution, the well-mixed model is a stochastic model. In order to run reproducible simulations, we need to specify the random number generator used by the stochastic sampler. We require that the cell cycle model is implemented using the `RandomState` Mersenne Twister pseudorandom number generator from the NumPy[4] random library. Moreover, the simulator optionally accepts a `seed` to initialise `RandomState` with. If no seed is given, it is initialised instead with a random seed supplied by the operating system.

##### S2.2.4 Termination conditions

The simulator class accepts simulation parameters that specify the simulation time and govern the termination conditions of the simulation. In particular, `tstart` and `tend` are the start and end times of the simulation, respectively, and `min_cell_count` and `max_cell_count` are the minimum and maximum cell counts, respectively. The simulation terminates when the simulation time reaches `tend`, when the cell count drops to `min_cell_count`, or when the cell count reaches `max_cell_count`; whichever event happens first. We summarise the simulation parameters in Table S1.

### S3 Implementation details of the vertex-based model

In this section, we describe the integration of the death clock framework into the vertex-based model using Chaste. As a versatile library for cell-based simulations, Chaste offers a high degree of customisation through abstract interfaces for modelling biological processes, such as apoptosis and intracellular reaction networks. Thus, we

[4] `https://numpy.org/`

| Name | Description | Type |
|---|---|---|
| `seed` | Seed for `RandomState` | integer |
| `tstart` | Simulation start time | float |
| `tend` | Simulation end time | float |
| `min_cell_count` | Minimum cell count | integer |
| `max_cell_count` | Maximum cell count | integer |
| `f_is_stepwise_constant` | Flag for stepwise constant death signal | Boolean |

Table S1: Summary of simulation parameters in the well-mixed model. The type refers to the data type used to store the parameter in the Python implementation.

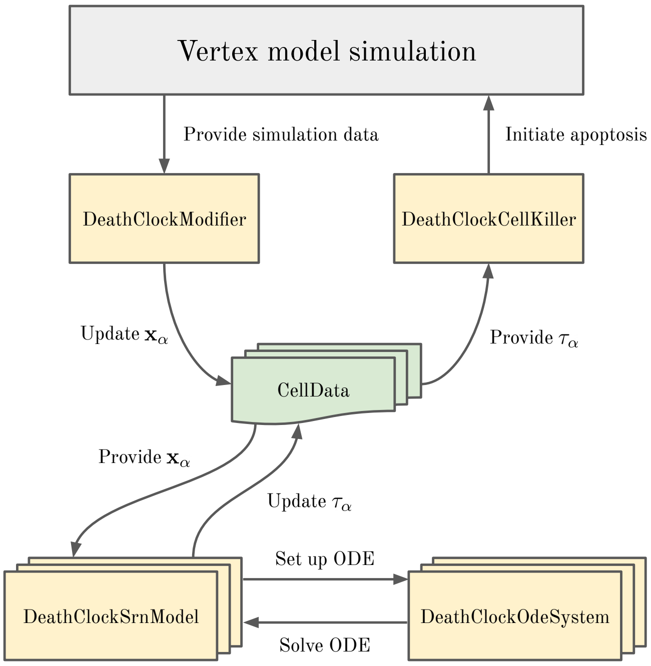


Figure S2: Diagram showing the components implementing the death clock framework in the vertex-based model and their interactions.

implemented the death clock framework by creating custom classes that adhere to these existing interfaces, rather than modifying the underlying codebase. The code for the vertex-based model with the death clock framework can be found in the following GitHub repository: `https://github.com/ThomasPak/cell-competition`.

#### S3.1 Overview

We store all the data related to the death clock framework in `CellData` objects, one for each cell. The `DeathClockModifier` extracts relevant data from the simulation to update the input vectors in `CellData`. These input vectors are fed to the `DeathClockSrnModel` of each cell to compute its death clock. Finally, the value of the death clock is read by the `DeathClockCellKiller` in order to enforce the apoptosis rule. See Figure S2 for a diagram summarising the interactions between these components.

#### S3.2 `CellData`

The `CellData` class specifies an object owned by each cell that holds an arbitrary amount of floating-point data. We use the `CellData` abstraction to store the cell state associated with the death clock framework. This includes the value of the death clock, death clock parameters such as the death threshold, as well as any simulation data that is relevant to the input vector.

#### S3.3 `DeathClockModifier`

The input vectors are updated by the `DeathClockModifier` class implementing Chaste's `SimulationModifier` interface. This is a general-purpose interface that provides read-and-write access to the entire simulation instance as it is being executed. For our purposes, the simulation modifier extracts simulation data relevant to the death clock at every timestep, specifically data that goes into the input vector, and stores them in `CellData` objects.

#### S3.4 `DeathClockSrnModel` and `DeathClockOdeSystem`

Chaste also offers functionality for modelling subcellular reaction networks with ODEs in tandem with a cell-based simulation. Once again, this is achieved through an object-oriented interface that lets the user specify a custom ODE system associated with each cell. The Chaste library also includes a suite of numerical ODE solvers.

We use this infrastructure to numerically integrate the death clock, i.e. Equation (20), on a per-cell basis. Concretely, we created the class `DeathClockOdeSystem`, which computes the right-hand side of Equation (20), and the class `DeathClockSrn-Model` (Srn is short for "subcellular reaction network"), which interfaces with both `DeathClockOdeSystem` and `CellData` to perform the logistical tasks involved with providing the ODE access to the input vector in `CellData`, running the numerical solver, and updating `CellData` with the computed death clock values.

#### S3.5 `DeathClockCellKiller`

Finally, we apply the apoptosis rule with the `DeathClockCellKiller` class using Chaste's abstractions for killing cells. Iterating over all cells at every timestep, the cell killer class accesses each cell's death clock, as well as its cell cycle phase, to determine whether the cell should undergo apoptosis. We note that the application of the apoptosis rule is procedurally decoupled from the numerical integration of the death clock, in contrast to the well-mixed model, because cell death and numerical integration are performed as distinct steps in the broader cell-based simulation scheme. As a result, the precise timing of apoptosis is constrained by the simulation timestep.

The vertex-based model also executes apoptosis differently than the well-mixed model. In the vertex-based model, cells enter an apoptotic state where they shrink in area until removed via a T2 swap when their area falls below a threshold. The exact timing of this removal is determined by the cell's mechanical interactions with its surroundings. Additionally, non-apoptotic cells can also be removed through

| Parameter | Well-mixed | Vertex-based |
|---|---|---|
| $t_G$ | | 100 |
| $c$ | | 1 |
| $\eta$ | | $0.05, 0.1, 0.2, 0.5, 1$ |
| $\beta$ | $0.1, 0.2, \ldots, 0.9$ | $0.2, 0.5, 0.8$ |
| Initial cell count | | 100 |
| Simulation end time | $\infty$ | 100 000 |
| Minimum cell count | | 10 |
| Maximum cell count | | 1 000 |
| $N_{\text{sim}}$ | 100 | 20 |

Table S2: Model and simulation parameter values used to estimate the homotypic survival probability.

"death by extrusion" which does not occur in the well-mixed model. Each T2 swap is recorded as either an "apoptosis" or "extrusion" event based on the cell's apoptotic state at the time of removal.

### S4 Computational validation of homotypic survival probability

This section compares the behaviour of the G2 death signal model in simulations to the homotypic survival probability predicted by Equation (31) using a Monte Carlo method. We ran multiple simulations with varying values of $\eta$ and $\beta$ to gather survival frequencies, and used a unique seed for the random number generator in each simulation to account for the model's stochastic nature.

#### S4.1 Parameter choice

We used the well-mixed model with parameters listed in Table S2 and chose $\eta$ values to obtain a range of survival probabilities above and below 1/2. For each $\eta$, we varied $\beta$ from 0.1 to 0.9 with increments of 0.1. We used the vertex-based model with parameters listed in Table S2 and limited our sample of $\beta$ values to 0.2, 0.5, and 0.8 due to computational constraints. The mechanical parameters of the vertex-based model were set to the default values specified in Table 1.

To translate the dimensionless parameters $\eta$ and $\beta$ to a concrete set of parameters, we fixed $t_G = 100$ and $c = 1$ and computed the death threshold as a function of the remaining parameters. In particular, rewriting Equation (30) gives

$$T_\dagger = \eta c t_G\,. \tag{S13}$$

#### S4.2 Initial conditions

In the well-mixed model, we must initialise the death clock, $\tau_\alpha$, the birth time, $t^0_\alpha$, and the G1 duration, $t^*_\alpha$, for every cell $\alpha$. For each cell, we set the death clock to zero and

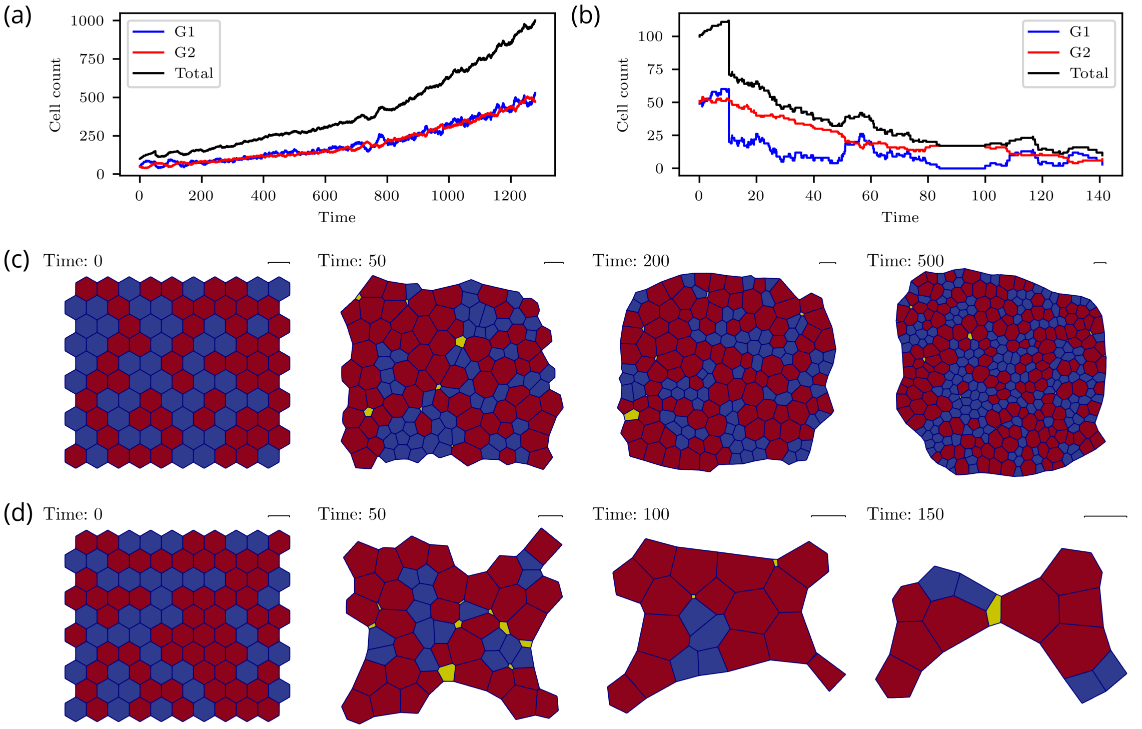


Figure S3: Example simulations of homotypic populations. (a–b) Cell counts for simulations of the well-mixed model. (a) $\eta = 0.2$, $\beta = 0.5$: the population proliferated until the simulation terminated by hitting the maximum cell count. (b) $\eta = 0.05$, $\beta = 0.5$: the population declined until the simulation terminated by hitting the minimum cell count. (c–d) Snapshots of simulations of the vertex-based model. Cells are coloured according to their state. Blue: G1 phase. Red: G2 phase. Yellow: apoptotic. (c) $\eta = 0.2$, $\beta = 0.5$: the population proliferated with few apoptotic cells until the maximum cell count was reached at $t = 1031$. (d) $\eta = 0.05$, $\beta = 0.5$: the population declined due to a high rate of apoptosis until the minimum cell count was reached at $t = 158$.

sampled the birth time uniformly from the interval $[-t_{G1} - t_{G2}, 0]$. Ideally, we would initialise the G1 duration by sampling directly from the exponential distribution, since this is how G1 durations are determined during simulation. However, naively sampling from the exponential distribution has the potential to clash with the previously assigned birth time by violating the division invariant. This happens when the total cell cycle duration, which is the sampled G1 duration plus fixed G2 duration, is less than the age of the cell, which is computed from its birth time. We avoided this conflict by iteratively sampling G1 durations until the division invariant was respected.

The initial conditions for the vertex-based model were determined in a similar manner, except for the G1 durations. Unlike the well-mixed model, the vertex-based model does not enforce a division invariant. Instead, Chaste checks at every timestep whether the cell age is greater than its cell cycle duration, and performs a division if so. The cell cycle duration is allowed to be smaller than the cell's age at the start of the simulation; it just means that the cell will divide as soon as the simulation starts. Hence, the G1 duration is sampled once from the exponential distribution for each cell. Finally, we arranged the cells in a honeycomb pattern for the initial spatial configuration, as shown in Figure S3(c–d).

#### S4.3 Termination conditions

We set minimum and maximum cell counts of 10 and 1 000, respectively. When the population hit either the minimum or maximum cell count, the simulation terminated. We set an unbounded simulation time for the well-mixed model, such that the simulation can only terminate by hitting the minimum or maximum cell count. On the other hand, there is no support in Chaste for an unbounded simulation time, so we set a simulation time of 100 000, which is large relative to the total cell cycle time of $t_G = 100$.

These termination conditions are illustrated in Figure S3, where we plotted example simulations of the well-mixed and vertex-based models. Figures S3(a) and S3(b) plot the cell counts of well-mixed simulations that terminated by hitting the maximum and minimum cell counts, respectively. Similarly, Figures S3(c) and S3(d) show snapshots of vertex-based simulations that terminated by hitting the maximum and minimum cell counts.

#### S4.4 Data processing and visualisation

For each simulation, we computed the survival frequency, denoted $\hat{\lambda}$, using Equation (1). For the well-mixed model, cells can only die through apoptosis, so all death events are apoptosis events. However, cells in the vertex-based model can be eliminated by a T2 swap despite not being in an apoptotic state. If a cell is apoptotic when it is extruded, we classify it as an apoptosis event, and as an "extrusion" event otherwise. We count both apoptosis and extrusion events as death events.

We plot the observed distribution of survival frequencies using box plots and compare with theoretical predictions in Figure S4(a) for the well-mixed model and Figure S4(b) for the vertex-based model.

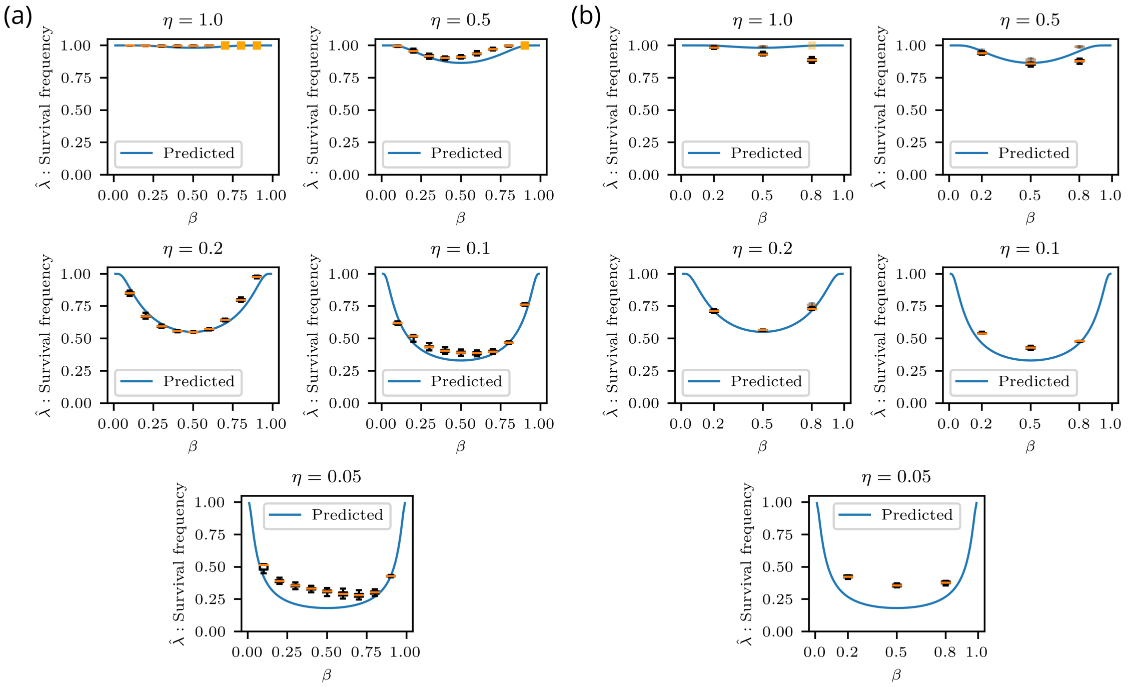


Figure S4: Homotypic survival probability. (a–b) Estimated homotypic survival frequency, $\hat{\lambda}$, defined in Equation (1), for the well-mixed and vertex-based models. The distribution of the survival frequency is plotted using box plots with whiskers extending to the minimum and maximum values. An orange square indicates that all observed survival frequencies are the same. The predicted homotypic survival probability, defined in Equation (31), is plotted for comparison. (a) Well-mixed model. (b) Vertex-based model when counting extrusions as death events (non-transparent), and without counting extrusions as death events (partially transparent).

#### S4.5 Results

Figure S4(a) generally shows good agreement between theory and observations for the well-mixed model, with minor deviations. The deviation from predictions is larger when the survival frequency is low, particularly below 1/2. This occurs when the rate of apoptosis is high and the limiting factor is no longer the survival probability, but the number of cells susceptible to apoptosis. In low-$\beta$ cell populations, cells spend less time in G1 phase, increasing their chance of evading apoptosis and resulting in a higher survival frequency than predicted. In high-$\beta$ cell populations, a larger fraction of cells are in G1 phase and for a longer time, increasing their vulnerability to apoptosis.

Figure S4(b) also shows good agreement between theory and observations for the vertex-based model. However, for high $\eta$ values, there are some discrepancies. For $\eta = 1.0$, the survival frequency is lower than expected and decreases with respect to $\beta$. This is due to a high survival frequency leading to a high proliferation rate, resulting in cell crowding and cell extrusion. Extrusion events are counted as death events, causing a lower survival frequency than expected from the death clock mechanism alone. This is shown in the partially transparent results in Figure 4(b), which

| Parameter | Well-mixed | Vertex-based |
|---|---|---|
| $t_{\mathrm{G}}$ | | 100 |
| $c$ | | 1 |
| $\eta$ | $0.01, 0.02, \ldots, 0.25$ | $0.02, 0.04, \ldots, 0.24$ |
| $\beta$ | $0.05, 0.10, \ldots, 0.95$ | $0.1, 0.2, \ldots, 0.9$ |
| Initial cell count | | 100 |
| Simulation end time | | 10 000 |
| Minimum cell count | | 10 |
| Maximum cell count | | 1 000 |
| $N_{\mathrm{sim}}$ | 50 | 20 |

Table S3: Model and simulation parameter values used to characterise the homotypic proliferation regimes.

demonstrate better agreement when extrusion events are excluded. Nonetheless, we still count extrusion events as death events because this effect only happens in cell crowding conditions and therefore does not affect the overall population viability.

### S5 Details for computational validation of homotypic proliferation regimes

In this section, we discuss details of the computational validation of homotypic proliferation regimes (Section 4.1.3). We used a Monte Carlo method, which involved running repeated well-mixed and vertex-based simulations for different values of $\eta$ and $\beta$, and sampling the homotypic survival frequency. For each simulation, we provided a unique seed to the random number generator.

We performed a systematic parameter sweep over a regular grid in $(\beta, \eta)$–space, spanning the area $\eta \in [0, 0.25]$ and $\beta \in [0, 1]$. The concrete parameter values used are listed in Table S3. We used a coarser grid for the vertex-based model than the well-mixed model because it is computationally more expensive. In addition, we used the default values for the mechanical parameters of the vertex-based model as specified in Table 1. We computed the death threshold by Equation (S13).

The initial conditions were determined in the same manner as in Section S4.2. We set minimum and maximum cell counts of 10 and 1 000, respectively. In addition, because the parameter sweep is relatively large, we set a simulation end time of 10 000 to cap the computational resources used by each simulation.

### S6 Computational validation of heterotypic survival difference

In this section, we verify whether simulations of heterotypic populations in the G2 death signal model match the predicted sign of the heterotypic survival difference in Equation (48). We used a Monte Carlo method to estimate the heterotypic

| Parameter | Cell type A | Cell type B |
|---|---|---|
| $t_{\mathrm{G}}$ | | 100 |
| $c$ | | 1 |
| $\eta$ | $0.01, 0.02, \ldots, 0.25$ | $0.05, 0.1, 0.2$ |
| $\beta$ | $0.05, 0.10, \ldots, 0.95$ | $0.3, 0.5, 0.7$ |
| Initial cell count | 50 | 50 |
| Simulation end time | | 10 000 |
| Minimum cell count | | 10 |
| Maximum cell count | | 1 000 |
| $N_{\mathrm{sim}}$ | | 50 |

Table S4: Model and simulation parameter values used to estimate the heterotypic survival difference for the well-mixed model.

| Parameter | Cell type A | Cell type B |
|---|---|---|
| $t_{\mathrm{G}}$ | | 100 |
| $c$ | | 1 |
| $\eta$ | $0.02, 0.04, \ldots, 0.24$ | 0.1 |
| $\beta$ | $0.1, 0.2, \ldots, 0.9$ | 0.5 |
| Initial cell count | 50 | 50 |
| Pattern | | random, segregated |
| Simulation end time | | 10 000 |
| Minimum cell count | | 10 |
| Maximum cell count | | 1 000 |
| $N_{\mathrm{sim}}$ | | 20 |

Table S5: Model and simulation parameter values used to estimate the heterotypic survival difference for the vertex-based model.

survival frequency of both cell types for different parameter values, and compared the difference in survival frequency with predictions. We provided a unique seed for the random number generator in each simulation.

#### S6.1 Parameter choice

Because there are two cell types in a heterotypic population, there are twice as many parameters compared to a homotypic population. Indeed, we want to verify our predictions in $(\beta_A, \eta_A, \beta_B, \eta_B)$–space. Because of the increase in dimensionality, we restricted the values of $\beta_B$ and $\eta_B$ to a limited set of cross sections, and varied $\beta_A$ and $\eta_A$ along a regular grid. For the well-mixed model, we picked $\beta_B = 0.3, 0.5, 0.7$, $\eta_B = 0.05, 0.1, 0.2$, evenly sampled 19 values for $\beta_A$ from the range $[0, 1]$, and evenly sampled 25 values for $\eta_A$ from the range $[0, 0.25]$. For the vertex-based model, we picked $\beta_B = 0.5, \eta_B = 0.1$, nine values for $\beta_A$, and 12 values for $\eta_A$. The mechanical parameters for both cell types are set to the default values specified in Table 1. See Tables S4 and S5 for a summary of the parameter values for the well-mixed and

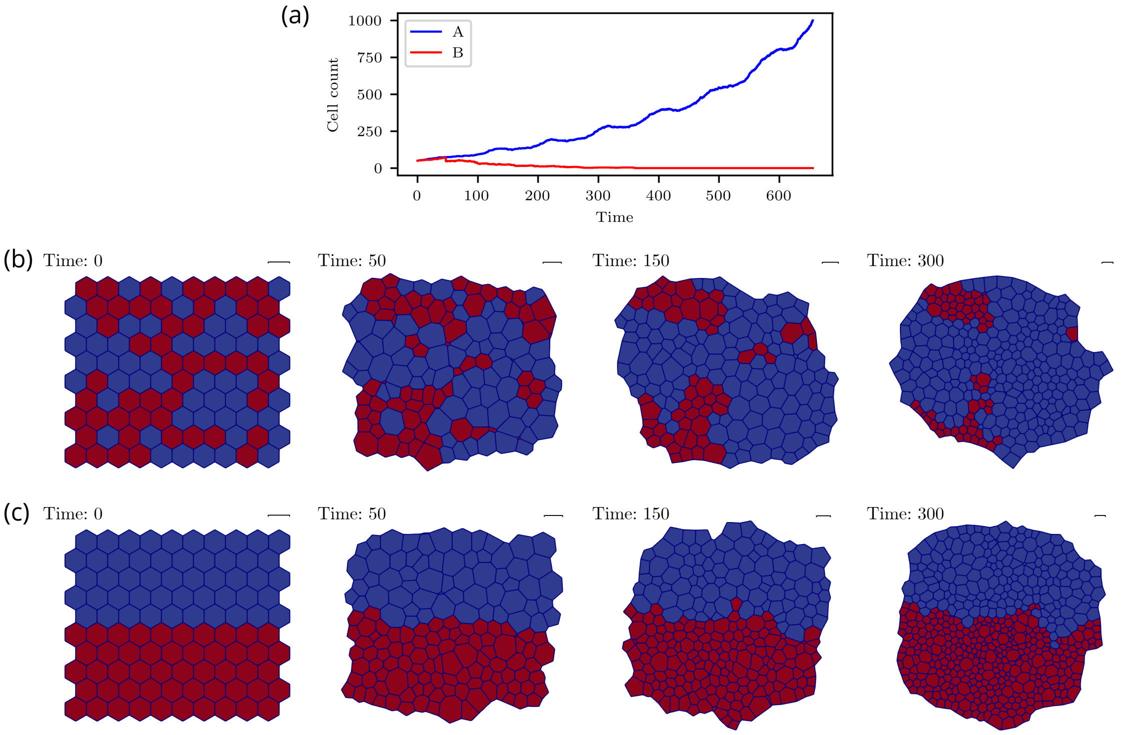


Figure S5: Example simulations of heterotypic populations. (a) Cell counts for a simulation of the well-mixed model with $\eta_A = \eta_B = 0.2$, $\beta_A = 0.2$, and $\beta_B = 0.8$. In well-mixed conditions, cell type A outcompetes cell type B. (b–c) Snapshots of simulations of the vertex-based model with $\eta_A = \eta_B = 0.2$, $\beta_A = 0.2$, and $\beta_B = 0.8$ for random and segregated initial conditions. Cells are coloured according to their cell type. Blue: cell type A. Red: cell type B. (b) Random: cell type A outcompetes cell type B. (c) Segregated: both cell types are viable.

vertex-based models, respectively.

To compute the death thresholds, we applied the approach in Section S4 to both cell types; we fixed $t_{\mathrm{G},A}, c_A, t_{\mathrm{G},B}, c_B$ and computed the death thresholds with the following expressions:

$$T_{\dagger,A} = \eta_A c_A t_{G,A}\,, \quad T_{\dagger,B} = \eta_B c_B t_{G,B}\,. \tag{S14}$$

#### S6.2 Initial conditions

In Section S4.2, we described the initial conditions for a homotypic population for the well-mixed and vertex-based models. We used the same initial conditions here, except that each cell was initialised in accordance with its cell type. In particular, the birth times and G1 durations were assigned using the cell cycle parameters $t_{\mathrm{G1},A}, t_{\mathrm{G2},A}$ for cell type A, and $t_{\mathrm{G1},B}, t_{\mathrm{G2},B}$ for cell type B. Furthermore, we initialised each simulation with 50 A-type cells and 50 B-type cells. See Figure S5(a) for an example simulation of the well-mixed model, terminating when the total cell count reaches 1 000.

For the vertex-based model we also needed to specify the initial spatial configuration. In order to test the effect of spatial segregation, we arranged the cell types either in a "segregated" or "random" pattern, similar to Section S1. To reiterate, in the former case, the cells were spatially segregated according to cell type, and in the latter case, cell types were distributed randomly in the tissue. See Figure S5(b–c) for examples of random and segregated initial conditions, respectively.

#### S6.3 Termination conditions

We set a minimum cell count of 10, a maximum cell count of 1 000, and a simulation end time of 10 000. We also terminated the simulation as soon as any cell type goes extinct, i.e. if either $n_A(t) = 0$ or $n_B(t) = 0$, since the population becomes homotypic in that moment. Hence, there are four different ways that the simulation can terminate: the simulation hits the minimum cell count, the simulation hits the maximum cell count, either cell type goes extinct, or the simulation runs until the end time.

#### S6.4 Data processing and visualisation

For each simulation, we computed the survival frequencies, $\hat{\xi}_{A|B}$ and $\hat{\xi}_{B|A}$, using Equations (3) and (4), respectively. Each parameter set was simulated $N_{\text{sim}}$ times with different random seeds. We used Equation (70) to estimate the survival frequency for a particular parameter set. The estimator for the heterotypic survival difference is then given by

$$\overline{\Delta^{\neq}_{A|B}} = \overline{\xi_{A|B}} - \overline{\xi_{B|A}} \,. \tag{S15}$$

We plot $\overline{\Delta^{\neq}_{A|B}}$ for fixed values of $\beta_B$ and $\eta_B$ using heat maps, with $\beta_A$ and $\eta_A$ on the horizontal and vertical axes, respectively. In addition, for the vertex-based model we decompose the results further into segregated and random initial conditions. We want to highlight the critical value $\overline{\Delta^{\neq}_{A|B}} = 0$ separating the regions where $\overline{\Delta^{\neq}_{A|B}} < 0$ and $\overline{\Delta^{\neq}_{A|B}} > 0$, respectively, so we use the diverging *seismic* colour map in Matplotlib[5]. Denoting the maximum absolute value of $\overline{\Delta^{\neq}_{A|B}}$ over all parameter sets as $|\Delta^{\neq}|$, we set the range of the colour map to $\left[-|\Delta^{\neq}|, |\Delta^{\neq}|\right]$ with central value zero. Hence, red regions and blue regions indicate positive and negative values, respectively.

We mark the degenerate case $\beta_A = \beta_B$, $\eta_A = \eta_B$ with a green dot and plot the coexistence curve with a solid line for comparison with theoretical predictions. The results are given in Figures S6 and S7 for the well-mixed and vertex-based models, respectively.

#### S6.5 Results

Figure S6 shows that the separation between $\overline{\Delta^{\neq}_{A|B}} > 0$ and $\overline{\Delta^{\neq}_{A|B}} < 0$ is generally consistent with the predicted coexistence curve for the well-mixed model. There are some deviations for $\eta_B = 0.05$ where the blue region does not tightly fit the

[5] `https://matplotlib.org/`

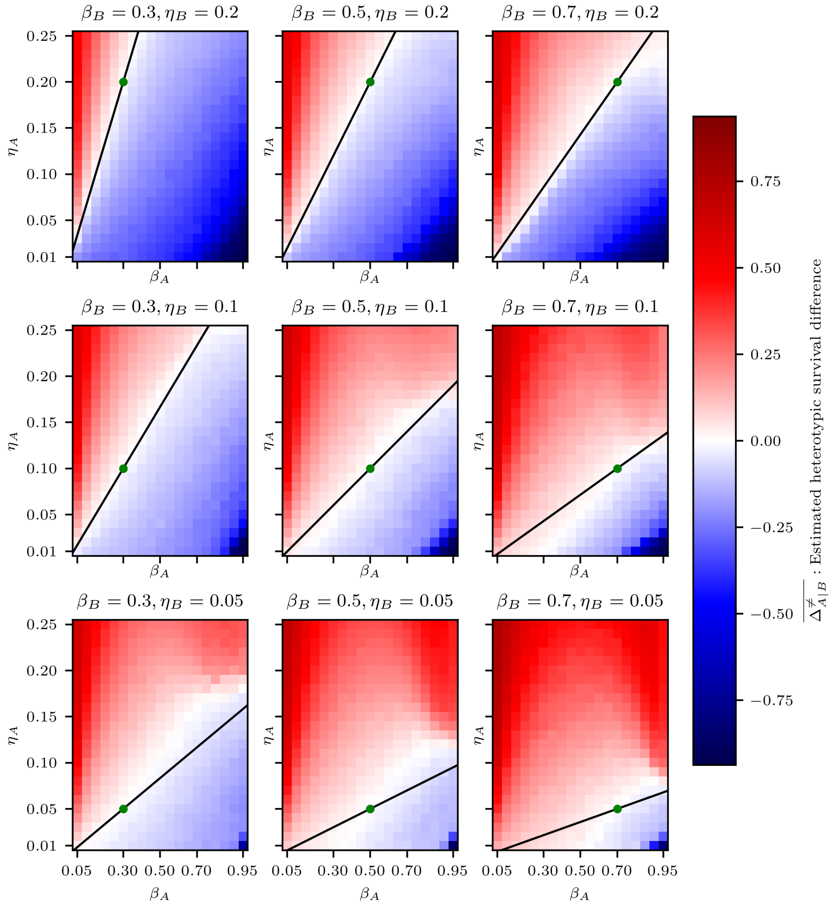


Figure S6: Estimated heterotypic survival difference, $\overline{\Delta^{\neq}_{A|B}}$, defined in Equation (S15), for the well-mixed model. The solid line and the green dot correspond to the coexistence curve and the neutral coexistence point, respectively.

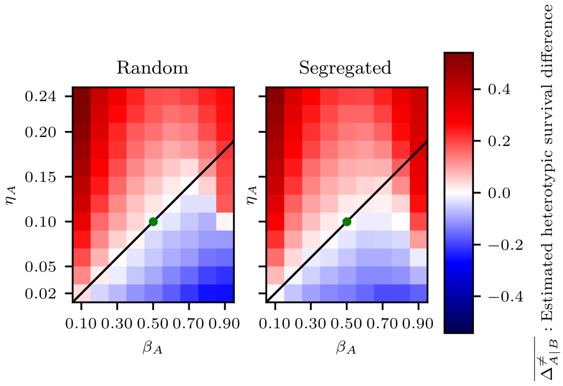


Figure S7: Estimated heterotypic survival difference, $\overline{\Delta^{\neq}_{A|B}}$, defined in Equation (S15), for the vertex-based model. The solid line and the green dot correspond to the coexistence curve and the neutral coexistence point, respectively.

coexistence curve. The overall structure of the parameter space remains intact, however.

Figure S7 shows that the observed parameter regions match the analytical predictions reasonably well for the vertex-based model. The largest deviation is found in the top right corner of the bottom triangular region, where the red region transgresses the predicted coexistence curve. Moreover, comparing the range of the colour bar with Figure S6 reveals that the observed values for $\overline{\Delta^{\neq}_{A|B}}$ are less extreme than for the well-mixed model. This effect is more pronounced for segregated initial conditions than for random initial conditions. In particular, the blue region in the bottom right corner is lighter. In addition, the red region under the coexistence curve is slightly larger for segregated simulations, suggesting that spatial segregation results in larger deviations from the well-mixed results.

### S7 Computational validation of homotypic survival difference

In this section, we verify whether the predicted sign of the homotypic survival difference in Equations (53) and (54) is supported by simulations. In order to estimate the homotypic survival difference, we need to run both homotypic and heterotypic simulations to estimate the survival probability in each, and combine the results. We have already run both types of simulations in Sections 4.1.3 and S6, respectively, so we combined the data from those sections to estimate the homotypic survival difference.

#### S7.1 Parameter choice

We used the parameters described in Section S5 and Section S6.1 to estimate homotypic and heterotypic survival probabilities, respectively. The parameters in those sections were chosen such that the values of $\beta_A, \eta_A, \beta_B, \eta_B$ in the heterotypic simulations coincide with the values of $\beta, \eta$ in the homotypic simulations.

#### S7.2 Data processing and visualisation

We reused the data from Section S6 for the heterotypic survival frequencies $\overline{\xi_{A|B}}$ and $\overline{\xi_{B|A}}$, and the data from Section 4.1.3 for the homotypic survival frequencies $\overline{\lambda_A}$ and $\overline{\lambda_B}$. We estimated the homotypic survival differences as follows:

$$\overline{\Delta^{=}_{A|B}} = \overline{\xi_{A|B}} - \overline{\lambda_A}\,, \quad \overline{\Delta^{=}_{B|A}} = \overline{\xi_{B|A}} - \overline{\lambda_B}\,. \tag{S16}$$

We plot $\overline{\Delta^{=}_{A|B}}$ and $\overline{\Delta^{=}_{B|A}}$ for fixed values of $\beta_B$ and $\eta_B$ using heat maps. We also split the results between segregated and random initial conditions for the vertex-based model. We denote the maximum absolute values of $\overline{\Delta^{=}_{A|B}}$ and $\overline{\Delta^{=}_{B|A}}$ across all parameter sets with $|\Delta^{\neq}_{A|B}|$ and $|\Delta^{\neq}_{B|A}|$, respectively. The colour map ranges are set to $\left[-|\Delta^{\neq}_{A|B}|, |\Delta^{\neq}_{A|B}|\right]$ and $\left[-|\Delta^{\neq}_{B|A}|, |\Delta^{\neq}_{B|A}|\right]$ with central values zero for the heat maps of $\overline{\Delta^{=}_{A|B}}$ and $\overline{\Delta^{=}_{B|A}}$, respectively. Hence, red and blue regions correspond to positive and negative values, respectively.

We mark the degenerate case $\beta_A = \beta_B$, $\eta_A = \eta_B$ with a green dot and plot the neutral competition curve with a dashed line for comparison with theoretical predictions. The well-mixed model results for $\overline{\Delta^{=}_{A|B}}$ and $\overline{\Delta^{=}_{B|A}}$ are given in Figures S8 and S9, respectively. The vertex-based model results for $\overline{\Delta^{=}_{A|B}}$ and $\overline{\Delta^{=}_{B|A}}$ are given in Figures S10(a) and S10(b), respectively.

#### S7.3 Results

Figures S8 and S9 show that the neutral curve in the parameter space is in good agreement with the border between red and blue regions for the well-mixed model. For $\overline{\Delta^{=}_{A|B}}$ in Figure S8, the parameter space is mostly dominated by blue regions where cell type A has a lower survival probability than in the homotypic case, whereas the red regions are relatively faint. This suggests that cell competition is driven more by the elimination of loser cells than the elevation of winner cells.

Moreover, the blue region has a characteristic triangular shape, transitioning into light blue towards the left and towards the top. To the left, the blue region overlaps with the Nonviable Regime of cell type A. Therefore, the A-type cells are homotypically nonviable to start with, and there is less room for B-type cells to further decrease their survival rate. To the top, the blue region is bounded by the coexistence curve. Above this curve, cell type A is the winning cell type, so it becomes the dominant species in the population and the effect of cell type B is diminished in comparison, leading to a smaller homotypic survival difference.

For $\overline{\Delta^{=}_{B|A}}$ in Figure S9, there are more pronounced red regions, particularly in the bottom right corner. This corresponds to the dark blue bottom right corner for $\overline{\Delta^{\neq}_{A|B}}$ in Figure S6. Inspection of individual heterotypic simulations reveals that A-type cells in this region perish virtually instantly so that B-type cells do not have the opportunity to initiate apoptosis before the A-type cells go extinct and the simulation terminates. As a result, the heterotypic survival frequency of B-type cells is recorded

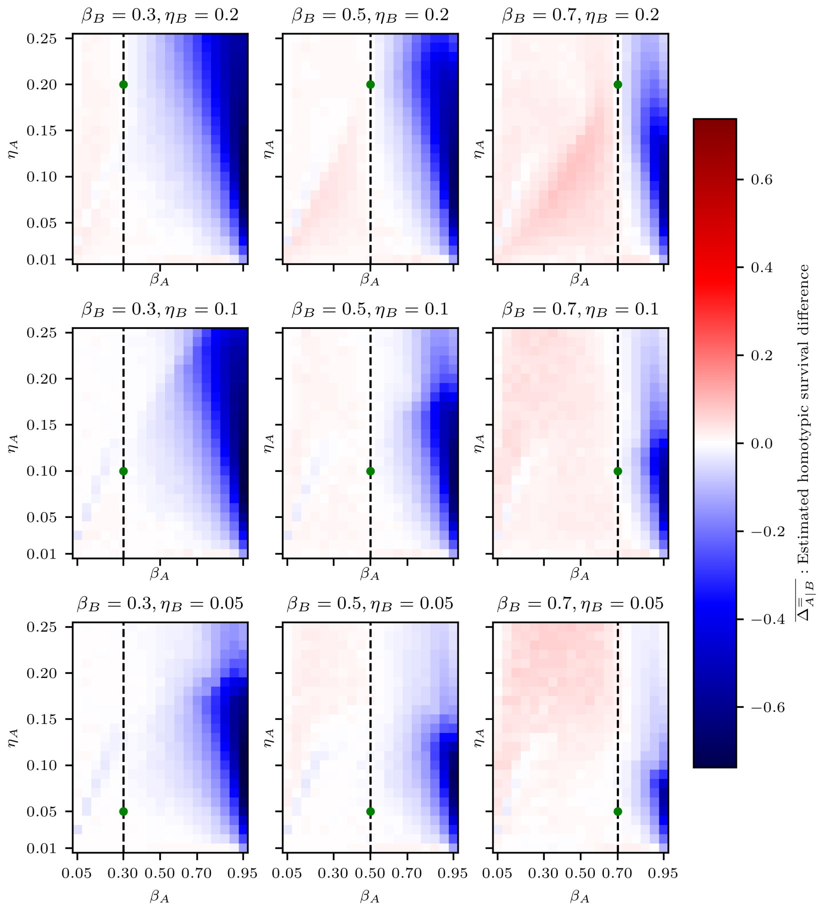


Figure S8: Estimated homotypic survival difference of cell type A, $\overline{\Delta^{=}_{A|B}}$, defined in Equation (S16), for the well-mixed model. The dashed line and the green dot correspond to the neutral competition curve and the neutral coexistence point, respectively.

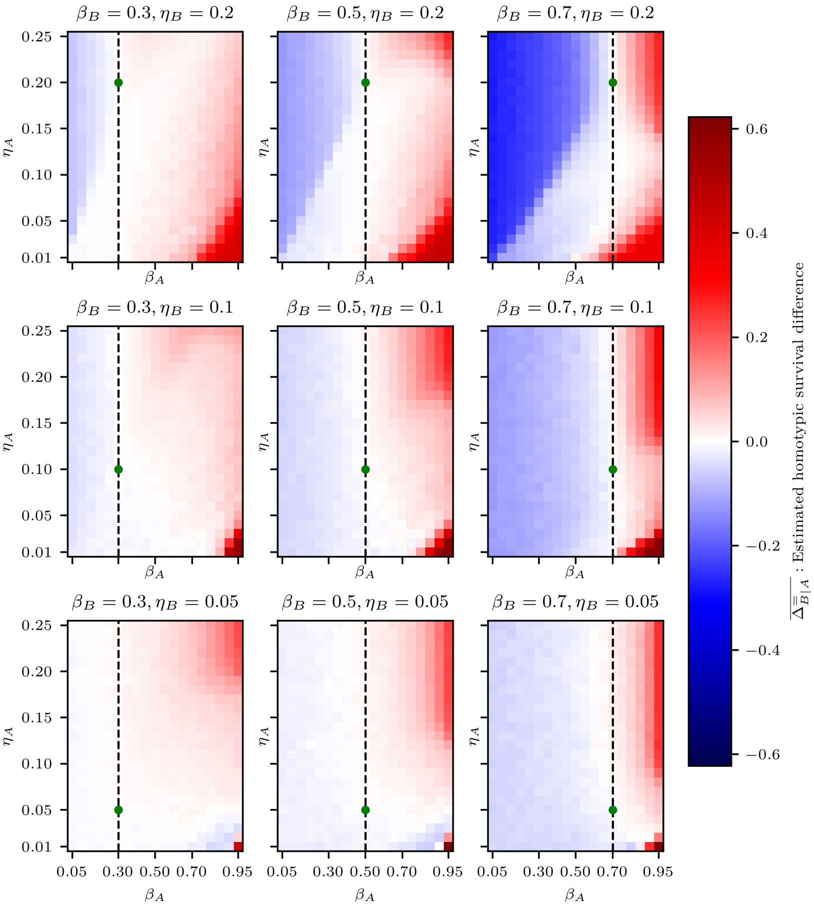


Figure S9: Estimated homotypic survival difference of cell type B, $\overline{\Delta^{=}_{B|A}}$, defined in Equation (S16), for the well-mixed model. The dashed line and the green dot correspond to the neutral competition curve and the neutral coexistence point, respectively.

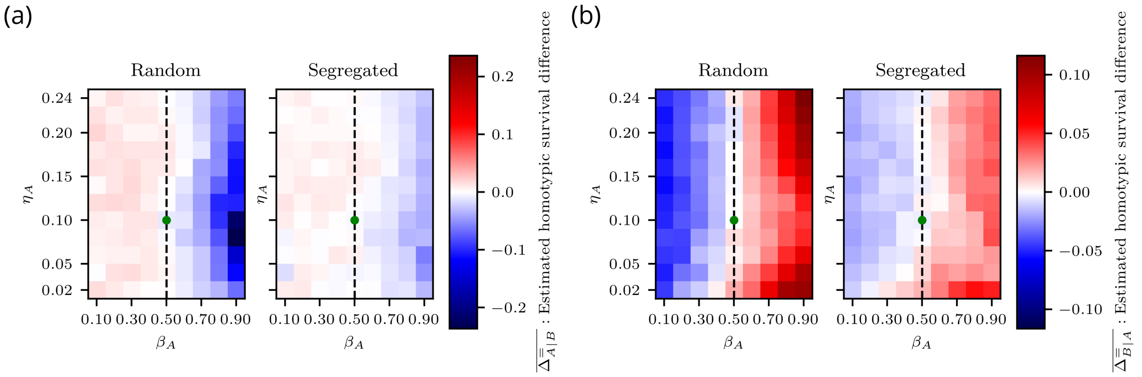


Figure S10: Estimated homotypic survival difference for vertex-based model with random and segregated initial conditions. The dashed line and the green dot correspond to the neutral competition curve and the neutral coexistence point, respectively. (a) Estimated homotypic survival difference of cell type A, $\overline{\Delta^{=}_{A|B}}$, defined in Equation (S16), for the vertex-based model. (b) Estimated homotypic survival difference of cell type B, $\overline{\Delta^{=}_{B|A}}$, defined in Equation (S16), for the vertex-based model.

as equal or close to unity, and we observe a high homotypic survival difference. We note, however, that this is a simulation artefact rather than a biologically meaningful phenomenon. Moving upwards from this corner, we encounter a faint middle region, followed by a dark upper region. Again, the border between these regions is roughly determined by the coexistence curve. Below the coexistence curve, cell type B is the winner, so the A-type cells are in the minority and have a limited impact on cell type B. Above this curve, A-type cells dominate and exert a larger influence on B-type cells.

In Figure S9, for $\beta_B = 0.7$, $\eta_B = 0.2$, a dark blue region is present in the top-left corner. This region is bounded on the right by the neutral curve and on the bottom by the coexistence curve. Above the coexistence curve, A-type cells are more abundant than B-type cells and therefore can exert their higher death signal to decrease the survival of B-type cells. Below the coexistence curve, B-type cells are more prevalent than A-type cells, thus limiting the influence of A-type cells as they are in lower numbers.

In Figures S10(a) and S10(b), it can be seen that the predicted neutral curve aligns well with the border obtained through simulation for the vertex-based model. The blue and red regions in these figures have similar shapes to those of the well-mixed model. However, when comparing the colour bars of these figures with those in Figures S8 and S9, it is apparent that the observed homotypic survival differences are smaller in magnitude than in the well-mixed case. Additionally, the figures indicate that this effect is more pronounced for segregated initial conditions. Given that $\overline{\Delta^{=}_{A|B}}$ and $\overline{\Delta^{=}_{B|A}}$ measure the impact of competition, this suggests that spatial segregation serves as a buffer against competition effects.

| Parameter | Cell type A | Cell type B |
|---|---|---|
| $t_{\mathrm{G}}$ | | 100 |
| $c$ | | 1 |
| $\eta$ | $0.01, 0.02, \ldots, 0.25$ | $\begin{bmatrix} 0.2 \\ 0.2 \end{bmatrix}, \begin{bmatrix} 0.2 \\ 0.8 \end{bmatrix}, \begin{bmatrix} 0.1 \\ 0.4 \end{bmatrix}$ |
| $\beta$ | $0.05, 0.10, \ldots, 0.95$ | |
| Initial cell count | 50 | 50 |
| Simulation end time | | 10 000 |
| Minimum cell count | | 10 |
| Maximum cell count | | 1 000 |
| $N_{\mathrm{sim}}$ | | 50 |

Table S6: Model and simulation parameter values used to estimate the heterotypic proliferation regimes for the well-mixed model.

| Parameter | Cell type A | Cell type B |
|---|---|---|
| $t_{\mathrm{G}}$ | | 100 |
| $c$ | | 1 |
| $\eta$ | $0.02, 0.04, \ldots, 0.24$ | $\begin{bmatrix} 0.2 \\ 0.2 \end{bmatrix}, \begin{bmatrix} 0.2 \\ 0.8 \end{bmatrix}, \begin{bmatrix} 0.1 \\ 0.4 \end{bmatrix}$ |
| $\beta$ | $0.1, 0.2, \ldots, 0.9$ | |
| Initial cell count | 50 | 50 |
| Pattern | | random, segregated |
| Simulation end time | | 10 000 |
| Minimum cell count | | 10 |
| Maximum cell count | | 1 000 |
| $N_{\mathrm{sim}}$ | | 20 |

Table S7: Model and simulation parameter values used to estimate the heterotypic proliferation regimes for the vertex-based model.

### S8 Details for computational validation of heterotypic proliferation regimes

In this section, we discuss details of the computational validation of heterotypic proliferation regimes (Section 4.2.6). We used a Monte Carlo method and ran multiple simulations to sample the survival frequency of cell types A and B for a wide range of parameter values. We used unique seeds for the random number generator in each simulation.

We restricted the values of $\beta_B$ and $\eta_B$ to the cross sections defined in Section 4.2.5. In each cross section, we used a regular grid in $(\beta_A, \eta_A)$–space to systematically investigate the parameter space. See Tables S6 and S7 for a summary of the parameter values for the well-mixed and vertex-based models, respectively. In addition, the mechanical parameters for both cell types in the vertex-based model are set to the default values given in Table 1. We computed the death thresholds for both cell types using Equation (S14). The initial conditions were determined in the same manner as in Section S6.2. We set a minimum cell count of 10, a maximum cell count of 1 000, and a simulation end time of 10 000.
